# Venous Angiogenesis and Dermal Fibroblast Augmentation of Decellularized Dermis: Towards Transplantable Dermal Substitutes

**DOI:** 10.64898/2026.09.30.755543

**Authors:** Sze Wai Lau, Yu Ling Zhang, Dongyue Xie, Nicholas Makhoul, Charlotte Jaloux, Jake Barralet

## Abstract

The ability to generate vascularized tissue constructs supplied by surgically accessible vessels remains a major limitation in reconstructive tissue engineering. Previous work demonstrated that positioning porous biomaterials around a vein can induce extensive vascular sprouting and generate a vascularized tissue volume without creation of an arteriovenous loop.

However, this phenomenon has principally been demonstrated using rigid calcium-phosphate scaffolds, limiting its direct relevance to soft-tissue reconstruction. Here, we investigated whether a flexible decellularized dermal matrix could similarly be vascularized around a single unmodified vein and whether autologous dermal fibroblasts could further enhance this response.

Decellularized rat dermis, with or without autologous dermal fibroblast seeding, was implanted either subcutaneously in the dorsal region or around the femoral vein for 8 weeks. Following Microfil perfusion, three-dimensional micro-CT demonstrated an extensive branching vascular network within perivenous constructs. Vessel volume relative to tissue volume was 0.206 ± 0.055% in subcutaneous constructs without fibroblasts (n = 11) and 0.598 ± 0.190% with fibroblasts (n = 9), compared with 6.131 ± 1.413% in perivenous constructs without fibroblasts (n = 10) and 7.329 ± 1.215% with fibroblasts (n = 10). Histology and immunohistochemistry similarly showed greater cellular repopulation and substantially higher CD31-and α-SMA-positive vessel densities following perivenous implantation. Integrated two-way ANOVA of the same 8-week data confirmed a dominant effect of implantation site across the vascular outcomes, while fibroblast seeding significantly affected vascular volume, dermal remodeling, and nuclear density. The effect of fibroblast seeding on vascular volume was site dependent, with a significant implantation site × fibroblast interaction.

At 8 weeks, the vascularized dermal construct was completely dissected free from the surrounding tissues while maintaining its connection to the femoral vein. A small superficial incision into the isolated dermis produced visible bleeding, supporting an acute functional connection of the intradermal vascular network to the host circulation while the femoral venous connection was preserved. After flap elevation, perivenous constructs retained substantially greater vascular volume and showed more complete surface healing and organized skin architecture than subcutaneous constructs. Fibroblast seeding accelerated wound closure, while its effect on post-elevation vascular volume depended on implantation site: VV/TV decreased in subcutaneous constructs but increased in perivenous constructs.

These findings show that single-vein vascularization can extend from rigid calcium-phosphate biomaterials to a compliant biological extracellular-matrix scaffold and may provide a simplified axial vascularization strategy without arterialization or incorporation of an arterial vessel.

## 1. Introduction

The generation of vascularized tissue volumes remains one of the central challenges in regenerative medicine and reconstructive surgery[1–4]. Although host capillaries can invade implanted biomaterials from surrounding tissues, the distance over which diffusion can maintain cell survival is limited[1, 5–7]. Consequently, vascularization arising only from the periphery becomes increasingly inadequate as the dimensions of an implanted construct increase[4, 8, 9]. Establishing a defined vascular axis within a regenerating tissue may therefore provide an important means of generating larger, surgically transferable tissue volumes[10, 11].

Several vascular configurations have been explored to establish a defined axial blood supply in soft-tissue constructs. Arterialized venous flaps demonstrated that tissue could be perfused through the venous system by directing arterial inflow into a vein, but this approach still requires arterialization and has shown variable survival and venous congestion[12–16]. In tissue-engineering models, Tanaka et al. first demonstrated that an AV shunt loop placed within artificial dermis could generate vascularized tissue capable of supporting a skin graft and subsequent flap elevation[17]. They subsequently compared AV shunt loops directly with AV bundles surrounded by artificial dermis. The AV bundles were examined using either a distal-ligation or a flow-through configuration, and both supported neovascularization. AV shunt loops produced a larger volume of newly formed vascularized tissue within the artificial dermis than either the distal-ligation or flow-through AV bundle configurations, whereas extensive capillary sprouting was particularly evident with the distally ligated AV bundle[18]. This AV-bundle concept was subsequently developed further as a vascular carrier for prefabricated engineered skin flaps[19].

Among these strategies, arteriovenous (AV) loops have been particularly extensively investigated within biomaterial scaffolds[10, 18, 20, 21]. The AV loop provides a high-flow vascular axis and can generate extensive vascular networks for the creation of transplantable tissue constructs[20, 22]. AV loops have also been combined with artificial dermis to generate vascularized constructs capable of supporting skin grafting and subsequent flap elevation, while intact AV bundles have been used to vascularize acellular dermal matrices for subsequent elevation as axial flaps[17, 23, 24]. Collagen-based matrices have also been used to generate axially vascularized soft-tissue flaps using AV loop models[25]. More recently, AV loops have been used to vascularize acellular dermal matrices in a large-animal model, generating scaffold-based flaps that could subsequently be microsurgically transferred as free flaps[26]. Collectively, these studies demonstrate the progression of vascularized dermal constructs from in situ prefabrication and flap elevation to free-tissue transfer. However, these approaches continue to rely on an arterial component, while AV loops additionally require microsurgical creation of an arteriovenous anastomosis[10, 20].

Previous work from our group demonstrated that extensive vascular sprouting could instead be induced by positioning a microporous calcium-phosphate scaffold around an unmodified vein[27, 28]. The resulting vascular network formed around and within the biomaterial without arterialization of the vein and reached a vascular volume comparable to that generated using an arteriovenous loop[28]. Importantly, the vascularized construct remained perfused following mobilization, demonstrating the potential of a single vein to act as the vascular axis of a newly generated tissue volume. However, this response was demonstrated predominantly using rigid ceramic scaffolds intended for hard-tissue regeneration, and the extent of vascularization was strongly influenced by scaffold material[27, 29].

Whether a single unmodified vein can similarly support vascularization of a compliant soft-tissue matrix remains unclear. This represents an important distinction from previous soft-tissue vascularization approaches, which have generally relied on arterialized venous flow, an arteriovenous loop, an arteriovenous bundle, or an intact arteriovenous pedicle[17, 18, 23, 26]. Decellularized dermis is particularly attractive for testing this concept because it provides a collagen-rich extracellular matrix that can be folded, sutured, contoured, and incorporated into soft-tissue defects while supporting host-cell infiltration and tissue remodeling[1, 30]. Demonstrating extensive vascularization of such a matrix using a vein alone would therefore extend the venosome concept from rigid biomaterials to a reconstructively relevant soft-tissue scaffold while eliminating the need to incorporate an artery[28, 29].

Autologous dermal fibroblasts may provide a complementary cellular strategy for enhancing vascularization of decellularized dermal matrices. Dermal fibroblasts support endothelial vessel formation through paracrine signaling and matrix remodeling[31]. Fibroblasts have been shown to promote blood vessel formation through paracrine signaling, including TIMP-1 secretion, while VEGF-expressing dermal fibroblasts enhanced angiogenesis and wound healing in vivo[32, 33].

In the present study, we investigated whether positioning decellularized dermis around a single femoral vein could induce vascularization of the scaffold and whether autologous dermal fibroblast seeding could further enhance this response. Identical unseeded and fibroblast-seeded dermal matrices were implanted either subcutaneously in the dorsal region or around the femoral vein and evaluated after 8 weeks using three-dimensional micro-computed tomography, histology, vascular immunohistochemistry and DiI fluorescence. We hypothesized that the femoral vein would provide an intrinsic vascular axis capable of generating substantially greater vascularization within decellularized dermis than conventional subcutaneous implantation. To our knowledge, this study represents the first demonstration of extensive vascularization of a decellularized dermal matrix using a single unmodified vein without arterialization, incorporation of an artery, or creation of an arteriovenous loop.

## 2. Materials and Methods

### 2.1. Preparation of decellularized dermis

Full-thickness dorsal skin was harvested from euthanized male Wistar rats. The skin was shaved, cleaned with 70% ethanol, and aseptically excised. Residual subcutaneous fat and muscle were carefully removed, and the tissue was stored overnight at 4 °C in phosphate-buffered saline (PBS; Gibco, 10010023) containing 1% antibiotic-antimycotic solution (Gibco, 15240062).

The skin was cut into 14 × 8 mm sections and rinsed twice with sterile water. Samples were immersed in 1 M saline for 24 h under continuous agitation on a laboratory rocker, followed by rinsing with sterile water and mechanical removal of the epidermis using forceps. The tissue was subsequently incubated with DNase I (0.2 mg/mL; Sigma-Aldrich, 11284932001) for 20 min and thoroughly rinsed with sterile water. The resulting decellularized dermal matrices were stored in 50% (v/v) glycerol (Fisher Chemical, 56815) in sterile PBS at 4 °C. Twenty-four hours before implantation, the matrices were removed from the glycerol solution, rinsed, and immersed in sterile PBS for rehydration until surgery. The decellularized dermal matrices were donor-derived allogeneic scaffolds; the term autologous refers only to fibroblasts isolated from the recipient animal assigned to a fibroblast-seeded group.

### 2.2. Animal model

In this study, 58 healthy male Wistar rats, aged 8-10 weeks and weighing 350-450 g, were used. All procedures were approved by the McGill University Animal Care Committee under animal use protocols AUP 8075 and 7662. Animals were maintained under standard housing conditions with a 12 h light-dark cycle at 20 ± 2 °C and provided food and water ad libitum.

Animals were allocated to four experimental groups for the 8-week vascularization study: a subcutaneous group without fibroblasts (n = 11), a subcutaneous group with autologous dermal fibroblasts (n = 9), a perivenous group without fibroblasts (n = 10), and a perivenous group with autologous dermal fibroblasts (n = 10). Additional animals were used for flap elevation experiments at week 8: five rats in each subcutaneous group and four rats in each perivenous group, with and without fibroblast seeding, respectively.

Sample size was estimated based on preliminary micro-CT vascular volume fraction (VV/TV) data from subcutaneous and perivenous constructs. Preliminary VV/TV values were 0.197 ± 0.024% for the subcutaneous group and 7.162 ± 0.950% for the perivenous group (n = 4 per group). Based on the anticipated difference between two independent groups, with a two-sided α of 0.05 and 90% power, the estimated minimum sample size was five animals per group. Experimental group sizes of 9-11 animals per group were used for the primary implantation study, exceeding the calculated minimum. Animals were randomly allocated to the experimental groups. Histological and immunohistochemical analyses were performed blinded to experimental group.

### 2.3. Isolation and culture of autologous dermal fibroblasts

Approximately 2 weeks before implantation, six 3-mm full-thickness dorsal skin biopsies were collected from animals assigned to fibroblast-seeded groups. Skin fragments were washed with 70% ethanol for 30 s, followed by three washes in PBS containing 2% antibiotic-antimycotic solution. Samples were incubated with trypsin-EDTA (Gibco, 25200072) at 37 °C for 5 min to facilitate epidermal removal, minced into approximately 1 × 1 mm pieces, and digested in 0.1% collagenase type IV (Gibco, 17104019) at 37 °C for 16 h. The resulting suspension was filtered through a 100 µm strainer and twice through 70 µm strainers, then centrifuged at 500 × g for 5 min. Cells were cultured in DMEM (Gibco, 11965092) containing 10% fetal bovine serum (Gibco, 12483020) and 1% antibiotic-antimycotic solution.

### 2.4. Fibroblast labelling and seeding

Autologous dermal fibroblasts were labeled with DiI (Invitrogen, D3911) before seeding. The seeding procedure was adapted from Vitacolonna et al.[34, 35]. Rehydrated dermis was submerged in 8 mL medium within a 50 mL syringe and subjected to five cycles of low-pressure degassing to remove trapped air. The matrix was then seeded with DiI-labeled fibroblasts at 4 x 105 cells/cm² and centrifuged at 300 × g for five cycles of 1 min. Seeded matrices were maintained overnight under static culture conditions and implanted the following day.

### 2.5. Implantation of decellularized dermal matrices

Preoperative analgesia consisted of sustained-release buprenorphine (1 mg/kg) and carprofen (5 mg/kg), administered 30 min before surgery. General anesthesia was induced and maintained with isoflurane.

For the subcutaneous groups, a 1.5 cm dorsal skin incision was made and a subcutaneous pocket was created by blunt dissection. A rehydrated decellularized dermal matrix, with or without autologous dermal fibroblasts according to group allocation, was placed within the pocket without additional fixation, and the incision was subsequently closed.

For the perivenous groups, a 1 cm incision was made along the inguinal fold and the underlying tissue was carefully dissected to expose the femoral vascular bundle. The femoral vein was isolated sufficiently to permit placement of the dermal matrix without interrupting blood flow. The decellularized dermis, with or without autologous dermal fibroblasts according to group allocation, was folded around the femoral vein and secured in position using two 4-0 Prolene sutures (Ethicon). The subcutaneous tissue and skin were subsequently closed using 5-0 Vicryl sutures (Ethicon).

Lidocaine was administered locally at the surgical site. Postoperative analgesia consisted of daily carprofen for 5 days, with an additional dose of sustained-release buprenorphine administered on postoperative day 3. To minimize self-trauma and wound dehiscence, an Elizabethan collar was applied immediately after surgery and maintained for approximately 7–10 days, until the incision had adequately healed. Animals were monitored regularly throughout the 8-week implantation period.

### 2.6. Intraoperative assessment of construct perfusion

At 8 weeks, the perivenous construct was exposed under anesthesia and carefully mobilized from the surrounding tissues while preserving continuity of the femoral vein. After mobilization, patency of the femoral vein was visually confirmed. A small superficial incision was then made in the dermal matrix with a scalpel, and the presence of bleeding from the construct was recorded. The procedure was documented in Supplementary Video S1.

### 2.7. Flap elevation and wound monitoring

Flap elevation was performed 8 weeks after implantation. Five animals from each subcutaneous group and four animals from each perivenous group underwent flap elevation. Subcutaneous constructs were exposed and mobilized, then folded and secured with approximately half of the dermal surface left exposed and sutured to the surrounding superficial skin. Perivenous constructs were similarly mobilized while maintaining continuity of the femoral vein before fixation to the surrounding skin.

Serial photographs were obtained after flap elevation to assess wound area, eschar formation, and re-epithelialization. Wound area was quantified from serial photographs and expressed as a percentage of the exposed area at the time of flap elevation. Complete wound closure was defined as the absence of a measurable exposed wound area.

### 2.8. Vascular perfusion and tissue harvesting

After 8 weeks of implantation, vascular perfusion was performed to visualize the newly formed vascular network. The thoracic cavity was opened to expose the heart, and approximately 200 mL of heparinized PBS containing heparin at a final concentration of 5 IU/mL (Sigma-Aldrich, H3149) was infused through the left ventricle using an 18G needle under gravity flow. A small incision was made in the right atrium to permit drainage. No paraformaldehyde perfusion was performed before contrast-agent injection. For the flap-elevation cohort, vascular perfusion and tissue harvesting were performed 4 weeks after flap elevation using the same procedure.

Yellow Microfil vascular contrast agent (MV-122, Flow Tech) was prepared by mixing 10 mL of yellow Microfil compound with 10 mL of diluent and 1 mL of curing agent. The contrast mixture was injected through a 20G catheter positioned in the descending abdominal aorta until adequate filling of the vascular circulation was observed. The descending aorta and inferior vena cava were then ligated, and the Microfil was allowed to polymerize overnight at 4 °C.

Following polymerization, the implants were carefully dissected from the surrounding tissues. For the subcutaneous group, the isolated dermal construct was harvested, whereas for the perivenous group, the dermal construct was harvested together with the enclosed femoral vein. Extraneous surrounding soft tissue was removed before subsequent analysis. Samples were fixed in 4% paraformaldehyde for 24 h, rinsed with PBS, and stored in 70% ethanol until further analysis.

### 2.9. Micro-computed tomography

Micro-computed tomography was used to visualize and quantify the three-dimensional vascular network within the implanted dermal constructs following Microfil perfusion. All available samples from both the 8-week vascularization cohort and the post-elevation cohort were analyzed by micro-CT. Samples were scanned using a SkyScan 1172 system (Bruker) with a 0.5 mm aluminum filter at an isotropic voxel size of 13 μm.

Projection images were reconstructed using NRecon software and analyzed using CTAn software (Bruker). Prior to scanning, surrounding tissues were removed so that the region of interest corresponded to the recovered dermal construct and its associated vasculature.

Vessel volume was quantified within the implanted dermal region and expressed relative to total tissue volume as vessel volume/tissue volume (VV/TV, %). In the perivenous group, the native femoral vein was segmented separately and its volume was subtracted from the total segmented vascular volume so that the reported vessel volume represented implant-associated perfused vasculature rather than the pre-existing axial vein.

### 2.10. Histology, immunohistochemistry and fluorescence imaging

Histological characterization was performed both before and after implantation. Prior to implantation, native rat skin and decellularized dermal matrices were processed for paraffin embedding and sectioned at a thickness of 4 µm. Adjacent sections were stained with hematoxylin and eosin (H&E) for histological assessment. Additional adjacent sections were stained with Masson’s trichrome using Bouin’s solution (Sigma-Aldrich, HT10132) and a Masson’s Trichrome Stain Kit (Sigma-Aldrich, HT15) to assess the preservation and organization of the collagen-rich extracellular matrix.

Following the 8-week implantation period, explanted constructs were paraffin embedded, sectioned at 4 µm, and stained with H&E to evaluate tissue morphology, host-tissue integration, cellular repopulation, structural remodeling, and vascular structures within the implanted dermal matrix. Six animals were randomly selected from each experimental group for immunohistochemical analysis.

For quantitative H&E analysis, the implanted dermal region was delineated as the total tissue area. Remodeled regions were defined by loss or reorganization of the original dense dermal collagen architecture together with host-tissue infiltration and were expressed as a percentage of total implant area. Nuclei within the implanted dermal region were counted and normalized to implant area (nuclei/mm²).

Immunohistochemistry was performed on serial sections to further characterize the vascular network within the implanted dermis. Following deparaffinization and rehydration, heat-induced antigen retrieval was performed using citrate buffer at pH 7. Endogenous peroxidase activity was blocked using hydrogen peroxide blocking reagent (Abcam, ab83781) for 10 min, followed by protein block (Abcam, ab81529) for 30 min.

Sections were incubated overnight at 4 °C with either a rabbit monoclonal anti-CD31 antibody (clone EPR17259, Abcam, ab182981; 1:1000) or a mouse monoclonal anti-α-smooth muscle actin (α-SMA) antibody (clone 1A4, Thermo Fisher Scientific, 14-9760-82; 1:250). For CD31 staining, sections were incubated with goat anti-rabbit HRP conjugate (Abcam, ab85999) for 15 min. For α-SMA staining, sections were incubated with mouse-specific reagent (Abcam, ab87648) for 10 min, followed by goat anti-rabbit HRP conjugate (Abcam, ab85999) for 15 min. Immunoreactivity was visualized using DAB substrate (Abcam, ab88385) prepared with 50× DAB chromogen (Abcam, ab86075) at a ratio of 10 µL chromogen to 500 µL substrate. Sections were counterstained with hematoxylin (Abcam, ab220365) and treated with bluing reagent (Abcam, ab85005).

Whole-slide images were acquired using an Aperio ScanScope XT slide scanner (Leica Biosystems). Quantitative analysis was performed using QuPath software, version 0.5.1 (64-bit). The implanted dermal region was defined as the region of interest, and CD31-positive and α-SMA-positive vascular structures were quantified and expressed as positive vessels per mm² of implant area. CD31 and α-SMA staining was performed on serial sections from the same specimens. DiI fluorescence microscopy was used in fibroblast-seeded constructs to assess cell retention and distribution within the dermal scaffold. For depth-profile analysis, the dermal thickness was divided into eight consecutive 150-µm regions from the superficial, former epidermal-facing surface toward the deep dermal surface. In seeded constructs, the deep surface corresponded to the original fibroblast-seeding surface and, in perivenous constructs, the vein-facing interface. Mean DiI fluorescence intensity was quantified within each region and expressed in arbitrary units (a.u.); four samples were analyzed per condition.

### 2.11. Statistical analysis

Data are presented as mean ± standard deviation (SD). Statistical analyses were performed using GraphPad Prism. For the 8-week pre-flap-elevation endpoint, subcutaneous and perivenous groups were compared separately within the unseeded and fibroblast-seeded cohorts using two-tailed unpaired t-tests with Welch’s correction. These comparisons were performed on the original, untransformed data. The same 8-week VV/TV, CD31-positive vessel density, α-SMA-positive vessel density, remodeled area, and nuclear density data were subsequently analyzed using two-way ANOVA to assess the overall effects of implantation site, fibroblast seeding, and their interaction, followed by uncorrected Fisher’s LSD tests for the indicated pairwise comparisons. For these two-way ANOVA analyses, remodeled area, α-SMA-positive vessel density, and VV/TV were log-transformed, whereas CD31-positive vessel density and nuclear density were analyzed on the original scale. Parametric-model assumptions were assessed by inspection of residual quantile-quantile (Q-Q) plots and

Brown-Forsythe tests for homogeneity of variance. Post-flap-elevation VV/TV data were log-transformed and analyzed using two-way ANOVA followed by uncorrected Fisher’s LSD tests for the indicated pairwise comparisons. DiI fluorescence profiles across scaffold depth were analyzed using two-way repeated-measures ANOVA, with depth as the repeated factor and experimental condition as the between-subject factor, followed by Tukey’s multiple comparisons test at individual depths. Serial wound-area measurements were analyzed using mixed-effects models, with time as the repeated factor and fibroblast seeding as the between-subject factor, followed by Šídák’s multiple comparisons test at individual time points. For outcomes log-transformed for two-way ANOVA, the ANOVA and associated pairwise p values were calculated from the transformed data, whereas descriptive statistics and graphical y-axes are presented on the original, untransformed scale. Statistical significance was defined as p ≤ 0.05.

## 3. Results

### 3.1. Histological characterization of native skin and decellularized dermis

H&E staining demonstrated effective removal of cellular and adnexal structures following decellularization while preserving the overall architecture of the dermal extracellular matrix (Figure 3). Native rat skin showed the expected organization of the epidermis and dermis, with abundant cellular nuclei, hair follicles, vascular structures, and other dermal appendages. In contrast, the decellularized dermis lacked an identifiable epidermal layer and contained no visible hair follicles, pre-existing vascular structures, or nuclei detectable by H&E. Masson’s trichrome staining further demonstrated preservation of the collagen-rich dermal extracellular matrix following decellularization, with maintenance of the characteristic collagen bundle architecture despite removal of native cellular and adnexal structures (Figure 3).

**Figure 1.**
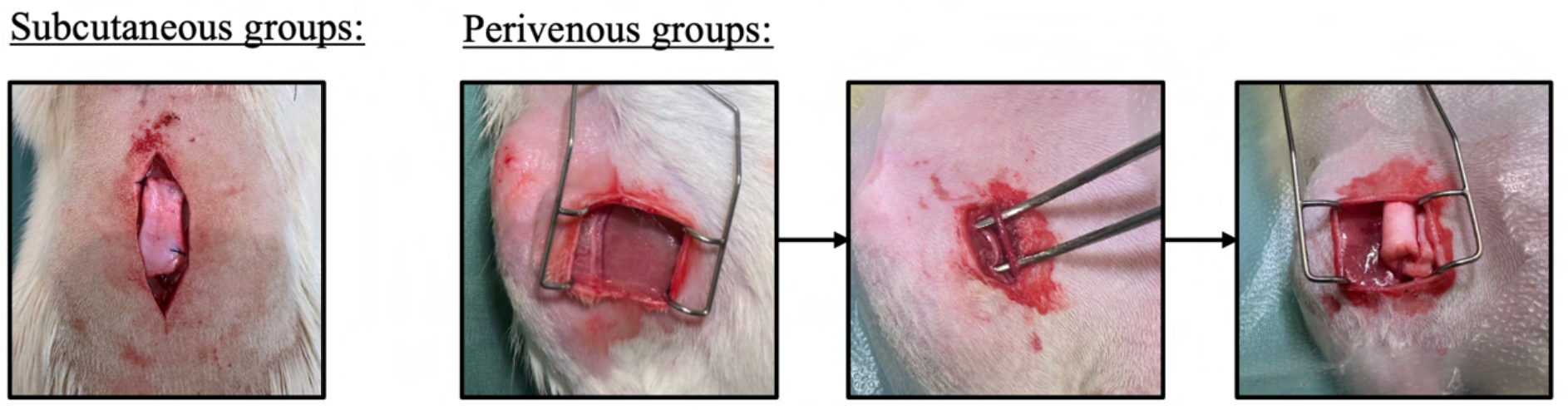
Surgical implantation of decellularized dermal matrices. Decellularized dermis, with or without autologous dermal fibroblast seeding, was implanted either within a dorsal subcutaneous pocket or folded around the femoral vein and secured with two sutures.

**Figure 2.**
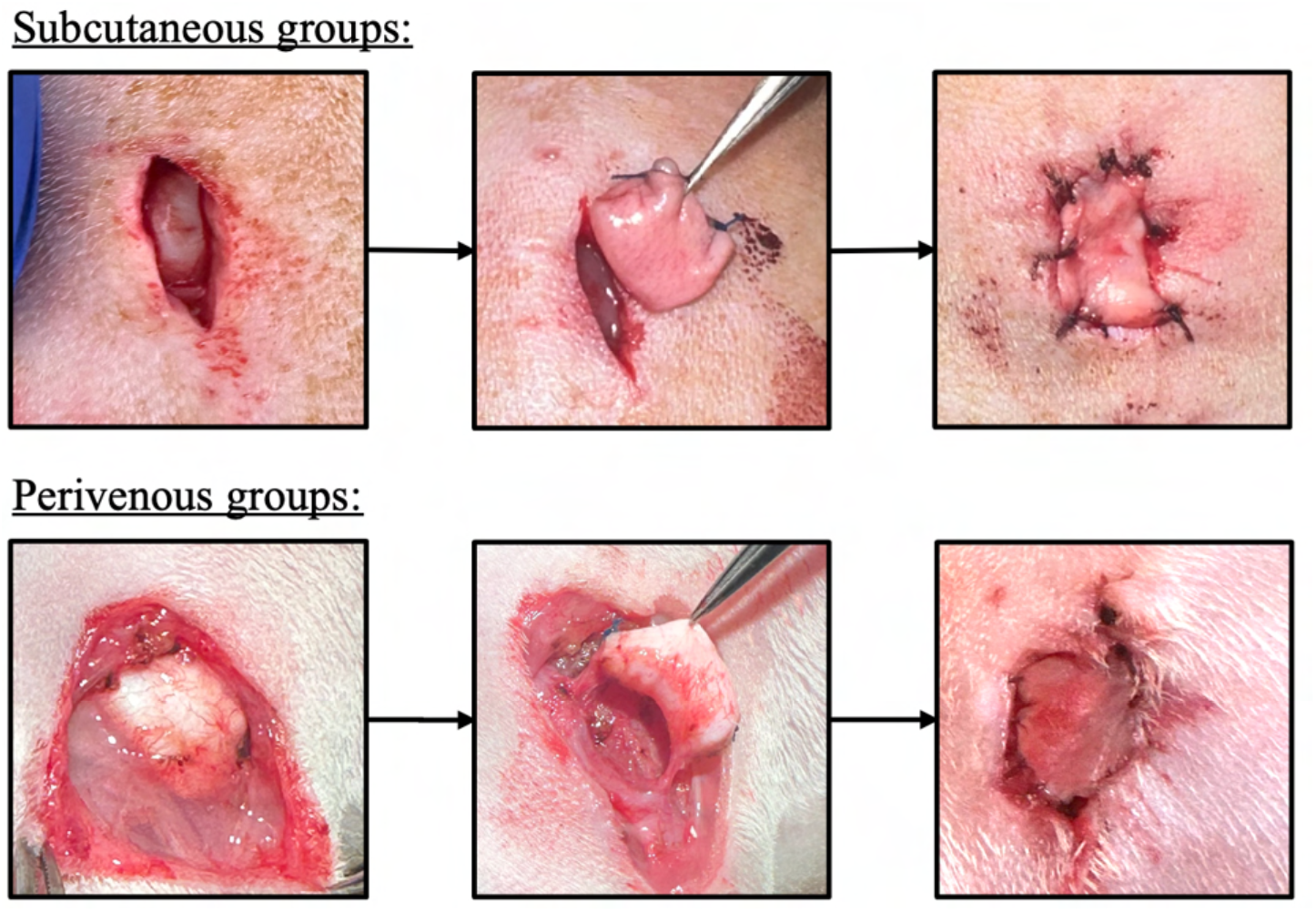
Flap-elevation procedure at week 8. The implanted dermal construct was mobilized and elevated 8 weeks after implantation. In the perivenous groups, the construct was mobilized while maintaining continuity with the femoral vein. The elevated construct was *secured to the surrounding skin with part of the dermal surface exposed for subsequent assessment of healing*.

**Figure 3.**
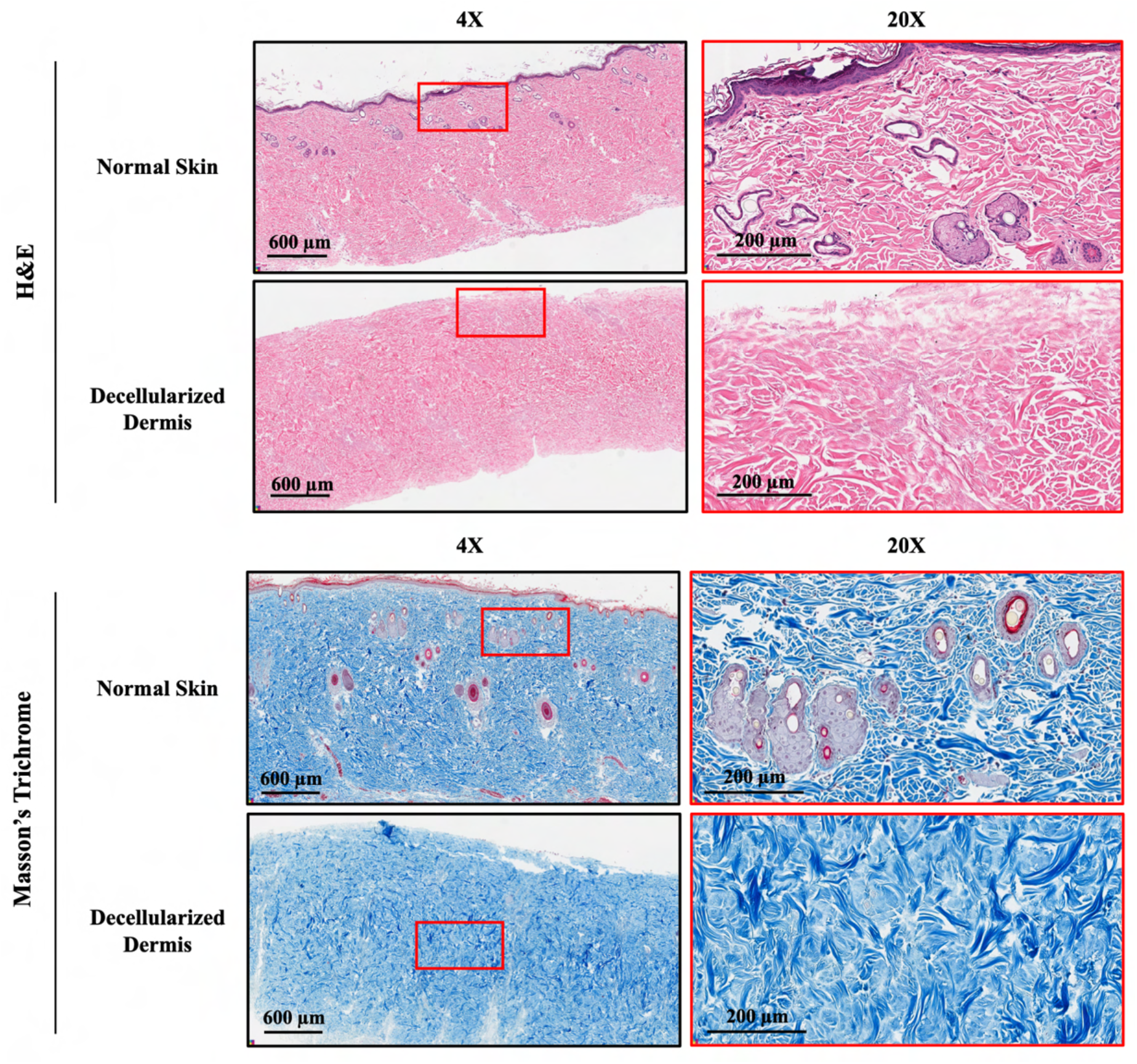
Histological characterization of native rat skin and decellularized dermis. Representative H&E and Masson’s trichrome sections of native skin and decellularized dermis. H&E staining demonstrates removal of cellular and adnexal structures following decellularization, while MT staining demonstrates preservation of the collagen-rich dermal extracellular matrix. Representative images are shown at 4× and 20× magnification. Scale bars = 600 µm at 4× and 200 µm at 20×.

### 3.2. Perivenous implantation markedly increased vascularization of unseeded decellularized dermis

Micro-CT following Microfil perfusion demonstrated a pronounced effect of implantation site on vascularization of unseeded decellularized dermis after 8 weeks (Figures 4 and 5). Subcutaneous constructs contained little detectable perfused vasculature within the dermal matrix, whereas perivenous constructs contained an extensive branching vascular network surrounding the femoral vein and extending throughout the matrix. Corresponding H&E sections supported these findings: vascular profiles and Microfil-containing vessels were sparse in subcutaneous constructs but abundant throughout the perivenous dermis (Figure 4).

**Figure 4.**
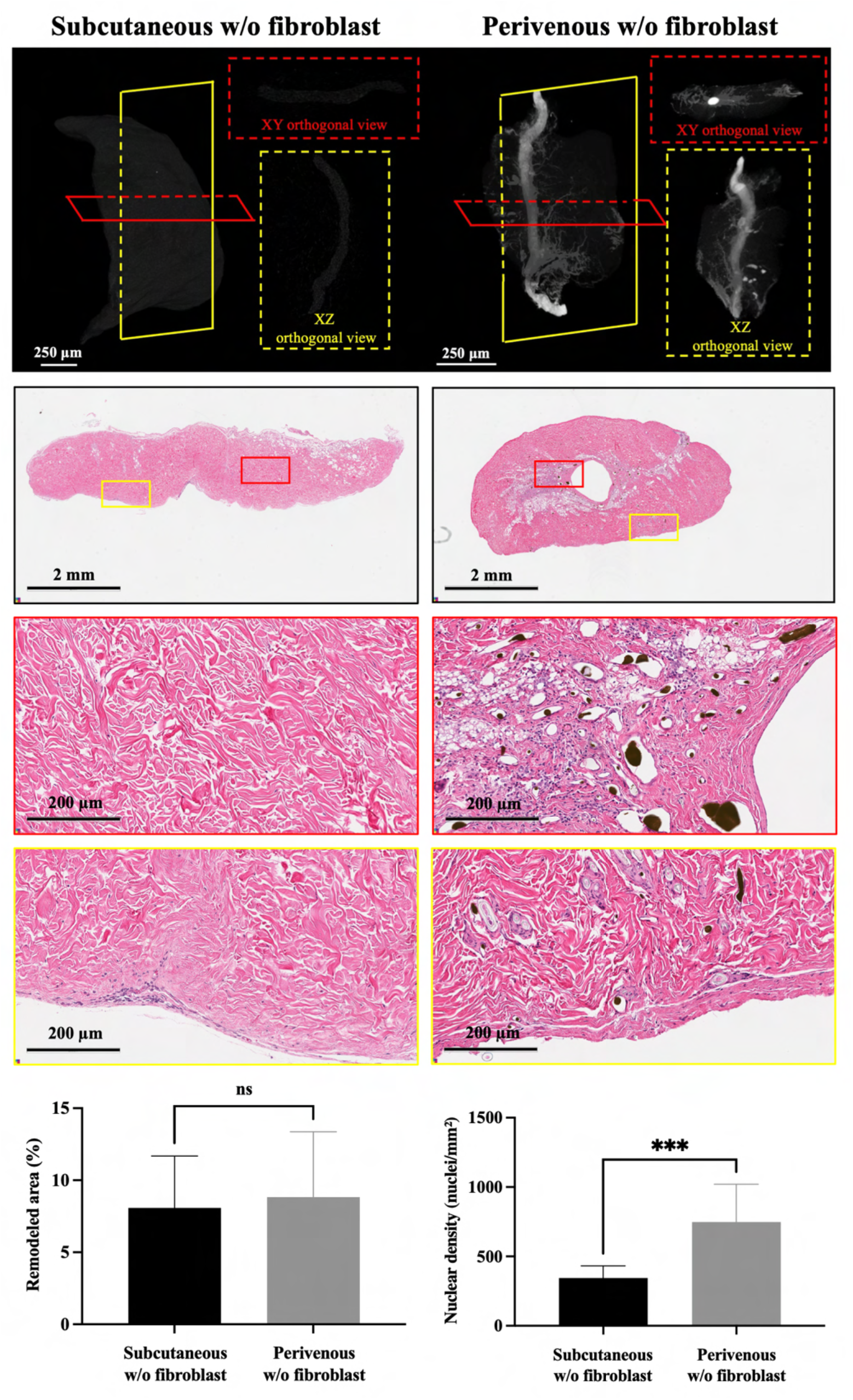
Three-dimensional vascularization and histological remodeling of unseeded decellularized dermal constructs after 8 weeks. Representative micro-CT reconstructions *following Microfil perfusion are shown for subcutaneous and perivenous constructs without fibroblast seeding. Red and yellow planes indicate the XY and XZ orthogonal planes, respectively. Representative H&E-stained sections show the overall morphology (2×) of the corresponding constructs, with boxed regions shown at higher magnification (20×). Microfil-containing vascular profiles were sparse in the subcutaneous construct and abundant throughout the perivenous dermal matrix. Remodeled area was quantified as a percentage of the total dermal construct area, and nuclear density was quantified as nuclei/mm². Data are presented as mean ± SD (subcutaneous, n = 11; perivenous, n = 10). Statistical analysis was performed using separate two-tailed unpaired t-with Welch’s correction. Scale bars are indicated in the individual panels*.

**Figure 5.**
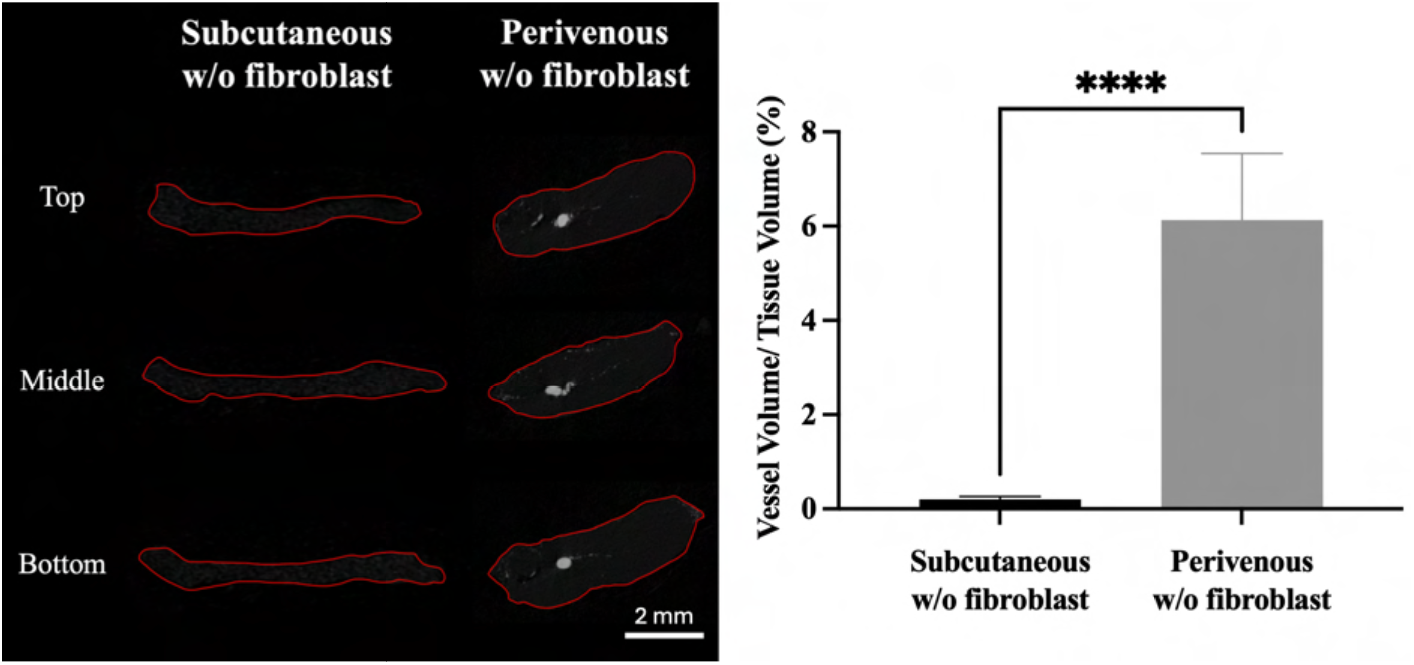
Perfused vascular volume in unseeded decellularized dermal constructs after 8 weeks. Representative transverse micro-CT sections from the top, middle, and bottom of subcutaneous and perivenous constructs without fibroblast seeding are shown, with the dermal implant outlined in red. Vessel volume relative to tissue volume (VV/TV, %) was quantified within the dermal implant. The native femoral vein was excluded from the vascular-volume calculation in the perivenous group. Data are presented as mean ± SD (subcutaneous, n = 11; perivenous, n = 10). Statistical analysis was performed using a two-tailed unpaired t-with Welch’s correction. ****p < 0.0001. Immunohistochemical assessment further confirmed the enhanced vascularization of the unseeded perivenous constructs (Figure 6). CD31-positive vessel density increased from 2.830 ± 0.929 vessels/mm² in the subcutaneous group to 33.67 ± 10.22 vessels/mm² in the perivenous group (p = 0.0007). Similarly, α-SMA-positive vessel density increased from 1.715 ± 0.494 vessels/mm² in the subcutaneous group to 26.54 ± 8.842 vessels/mm² in the perivenous group (p = 0.0010). Both comparisons were performed using two-tailed unpaired t-with Welch’s correction (n = 6 per group). These findings were consistent with the micro-CT results and demonstrated markedly greater endothelial and α-SMA-positive vascular structures following perivenous implantation.

**Figure 6.**
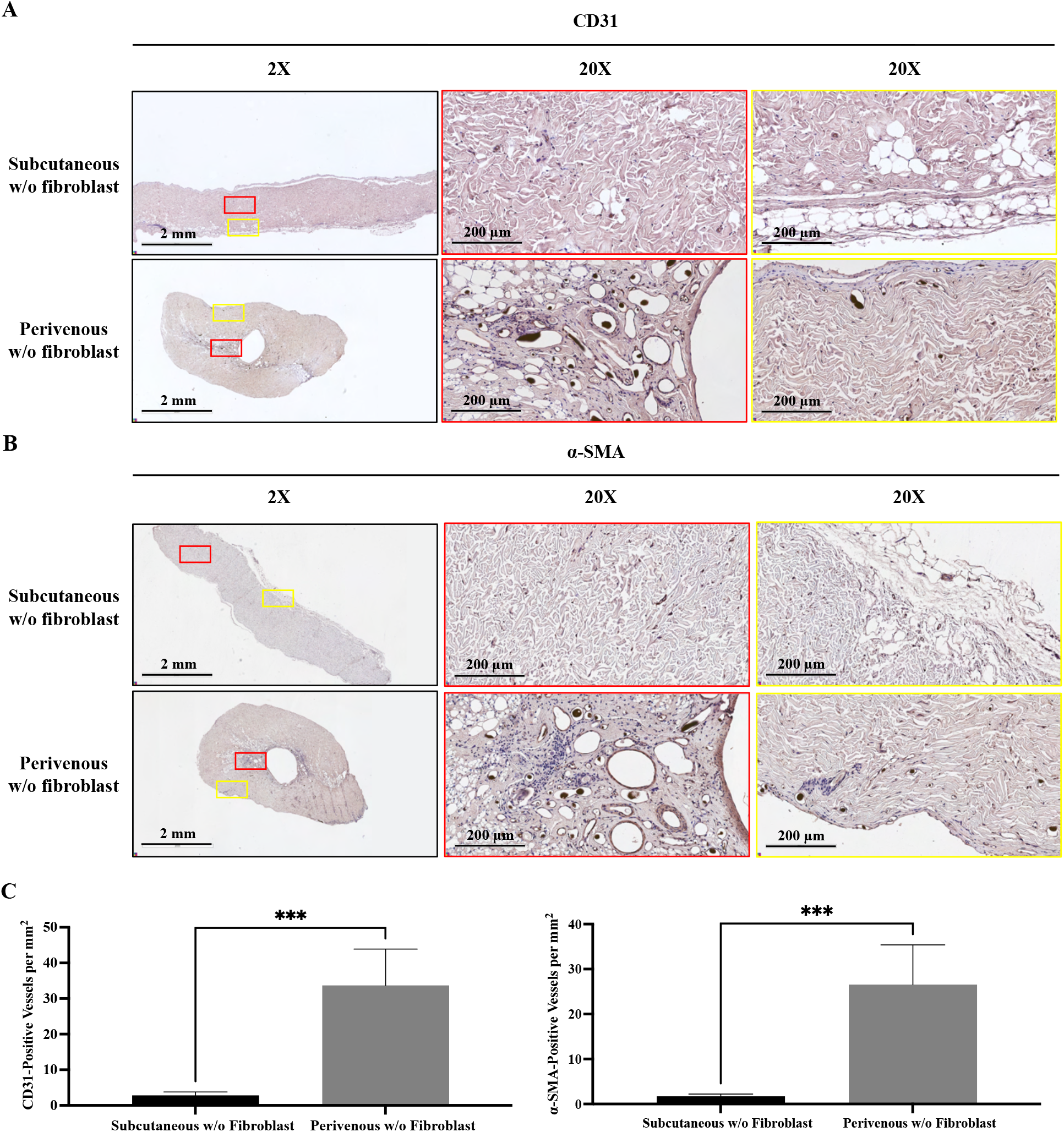
CD31-and α-SMA-positive vascularization of unseeded decellularized dermal constructs after 8 weeks. (A) Representative CD31 immunohistochemical staining of subcutaneous and perivenous constructs without fibroblast seeding at 2× and 20× magnification. (B) Representative α-SMA immunohistochemical staining of the corresponding groups at 2× and 20× magnification. Red boxes in the low-magnification images indicate the corresponding regions shown at higher magnification. (C) Quantification of CD31-positive *and α-SMA-positive vessel densities. Data are presented as mean ± SD (n = 6 per group). Statistical analysis was performed using separate two-tailed unpaired t-with Welch’s correction. ***p < 0.001. Scale bars: 2 mm at 2× and 200 µm at 20×*.

Beyond vascular morphology, H&E analysis was used to assess cellular repopulation and structural remodeling of the implanted dermal matrix. The proportion of remodeled dermal area was comparable between unseeded subcutaneous and perivenous constructs (8.09 ± 3.59% and 8.83 ± 4.54%, respectively; p = 0.6869). In contrast, nuclear density was markedly greater in perivenous constructs, increasing from 344.4 ± 87.7 nuclei/mm² in the subcutaneous group to 748.5 ± 272.7 nuclei/mm² in the perivenous group (p = 0.0010). Both comparisons were performed using two-tailed unpaired t-with Welch’s correction. These findings indicate that perivenous implantation substantially increased cellular repopulation of the unseeded dermal matrix without a corresponding increase in the proportion of histologically remodeled matrix at 8 weeks.

Consistent with the three-dimensional reconstructions, quantitative micro-CT confirmed the marked vascular advantage of perivenous implantation in the absence of fibroblast seeding (Figure 5). VV/TV increased from 0.206 ± 0.055% in subcutaneous constructs to 6.131 ± 1.413% in perivenous constructs (p < 0.0001; two-tailed unpaired t-test with Welch’s correction). The native femoral vein was excluded from the vascular-volume calculation so that the reported VV/TV represented implant-associated perfused vasculature rather than the pre-existing axial vein.

### 3.3. Fibroblast retention and distribution within decellularized dermis

Having established the marked site-dependent vascular response in unseeded constructs, we next assessed the retention and spatial distribution of the autologous dermal fibroblasts used in the seeded cohorts. After 7-10 days of isolation and in vitro expansion, autologous dermal fibroblasts were labelled with DiI before seeding onto decellularized dermis. The labelled cells showed bright fluorescence before seeding (Figure 7A, B), and DiI signal was readily detected within the scaffold immediately after seeding (Figure 7C). After 8 weeks, DiI-associated fluorescence remained detectable in both the in vitro and in vivo constructs, although its distribution through the scaffold differed markedly between conditions (Figure 7D-H).

**Figure 7.**
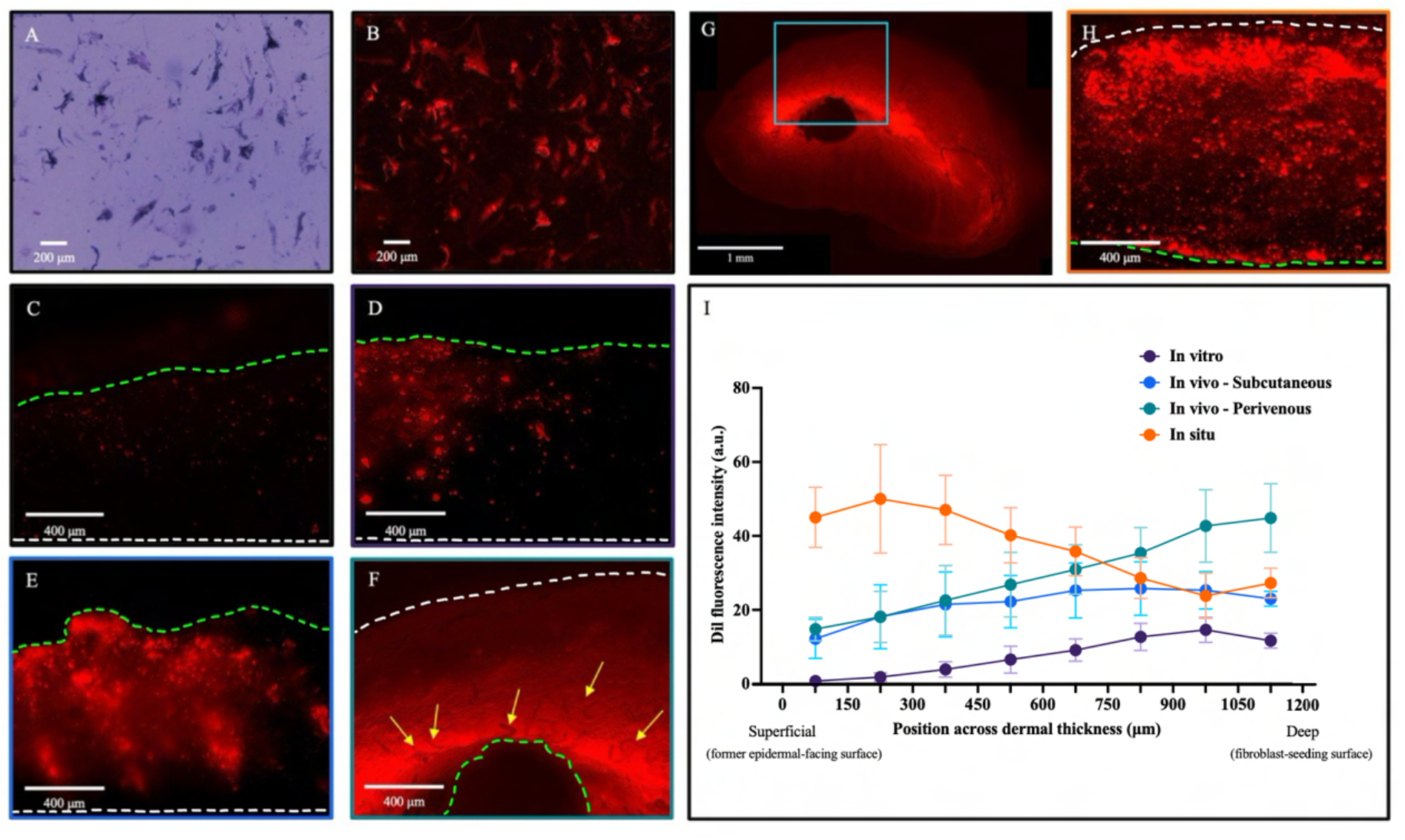
Distribution of DiI-associated autologous dermal fibroblasts in decellularized dermis. (A, B) Representative bright-field and fluorescence images of DiI-associated autologous dermal fibroblasts before seeding. (C, D) Representative fluorescence images of fibroblast-seeded decellularized dermis maintained in vitro at day 0 (C) and week 8 (D). (E) Representative fibroblast-seeded subcutaneous construct after 8 weeks in vivo. (F, G) Representative fibroblast-seeded perivenous construct after 8 weeks in vivo (F) and photomerged overview showing DiI-associated fibroblasts within the dermal matrix in relation to adjacent Microfil-perfused vessels (G). Yellow arrows in (F) indicate the femoral vein. (H) Representative fluorescence image of cells labelled in situ in native dermis. In (C–F, H), white and green dotted lines indicate the superficial (epidermal-facing) and deep dermal surfaces, respectively; in (C–F), the deep surface corresponds to the fibroblast-seeding surface. (I) Quantification of DiI fluorescence intensity across eight consecutive 150-µm regions from the superficial to deep dermal surface. Data are presented as mean ± SD (n = 4 per condition). Scale bars: A–B, 200 µm; C–F and H, 400 µm; G, 1 mm.

Quantification across eight consecutive 150-µm regions demonstrated distinct spatial fluorescence profiles across the dermal thickness (Figure 7I). Measurements were performed from the superficial, former epidermal-facing surface toward the deep dermal surface. In fibroblast-seeded constructs, the deep dermal surface corresponded to the original fibroblast-seeding surface and, in perivenous constructs, the vein-facing interface. DiI fluorescence ranged from 0.76 to 14.63 a.u. in vitro, 12.19 to 25.80 a.u. in subcutaneous constructs, 14.84 to 44.86 a.u. in perivenous constructs, and 23.81 to 50.03 a.u. in native dermis labelled in situ. In vitro constructs showed relatively little fluorescence in regions distant from the original seeding surface, whereas subcutaneous constructs showed an intermediate distribution. Perivenous constructs exhibited greater fluorescence extending throughout the dermal matrix, including regions distant from the original seeding surface. Native dermis labelled in situ showed a heterogeneous fluorescence distribution across the dermal thickness and was included as a qualitative reference for the spatial distribution of labelled cells in intact dermal tissue. Overall, perivenous implantation was associated with a broader distribution of fibroblast-associated DiI fluorescence through the dermal matrix than either in vitro or subcutaneous implantation.

Fluorescence imaging of the perivenous constructs further showed DiI-associated fibroblasts within the dermal matrix in close spatial association with the vascularized region (Figure 7F, G). The femoral vein was identifiable within the construct, with numerous adjacent Microfil-perfused vascular branches extending through the surrounding dermis (yellow arrows), while the photomerged overview demonstrated DiI-associated fibroblasts distributed within the same vein-associated matrix. Together with the depth analysis, these observations show that fibroblast-associated DiI fluorescence remained detectable after 8 weeks and was distributed extensively throughout the perivenous dermal construct.

### 3.4. Perivenous implantation maintained its vascular advantage in fibroblast-seeded constructs

Having confirmed the persistence and distribution of the seeded fibroblast-associated DiI signal, we next examined whether the site-dependent vascular response observed in unseeded constructs was maintained after fibroblast seeding and whether fibroblast addition provided further vascular enhancement. Micro-CT also demonstrated sparse perfused vessels in subcutaneous constructs and an extensive branching network throughout perivenous constructs (Figure 8). H&E sections similarly showed few vascular profiles in subcutaneous dermis but numerous Microfil-containing vessels within the perivenous matrix.

**Figure 8.**
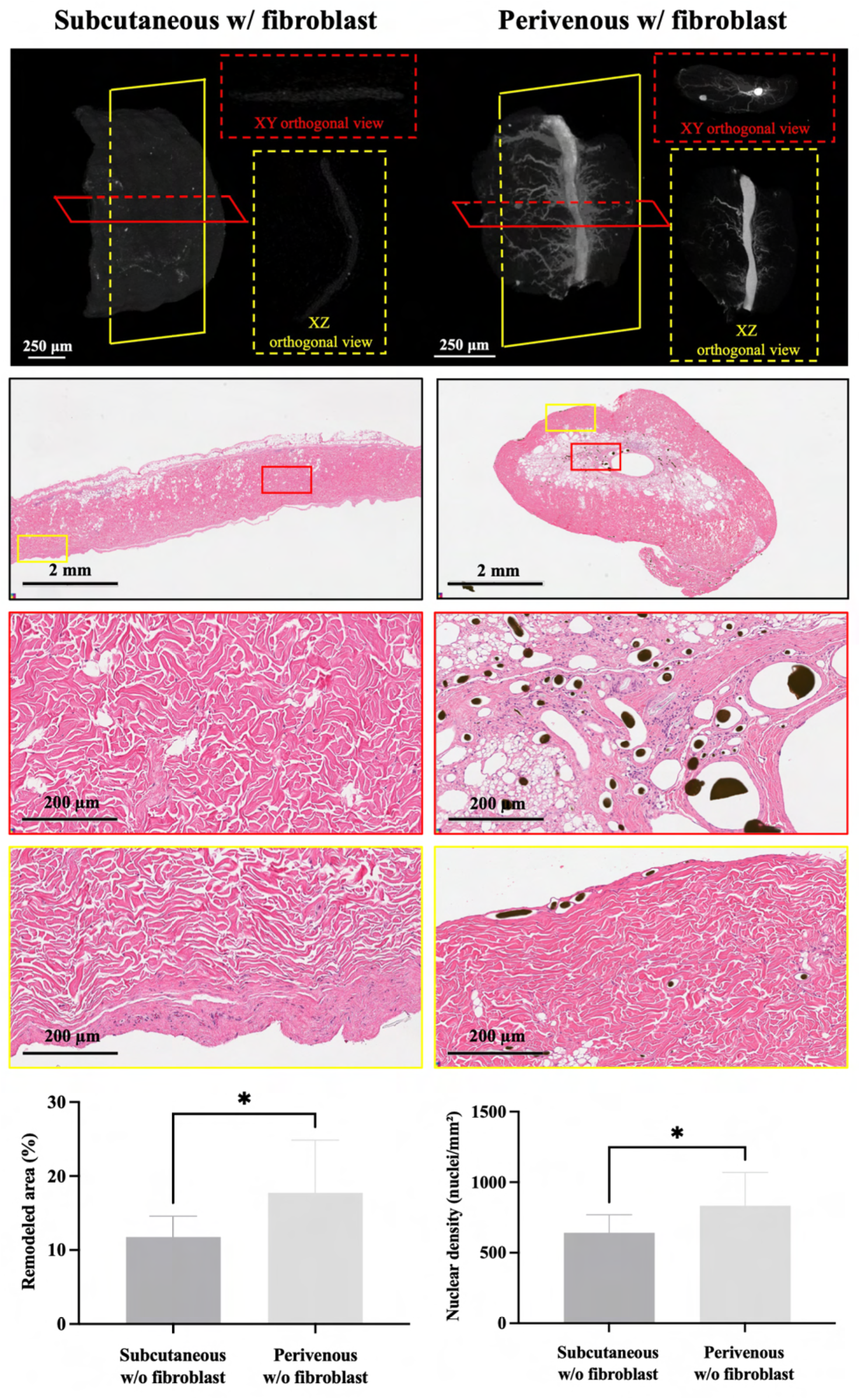
Three-dimensional vascularization and histological remodeling of fibroblast-seeded decellularized dermal constructs after 8 weeks. Representative micro-CT reconstructions following Microfil perfusion are shown for fibroblast-seeded subcutaneous and perivenous *constructs. Red and yellow planes indicate the XY and XZ orthogonal planes, respectively, with the corresponding orthogonal views shown adjacent to each three-dimensional reconstruction. Representative H&E-stained sections show the overall morphology (2×) of the corresponding constructs, with boxed regions shown at higher magnification (20×). Microfil-containing vascular profiles were sparse in the subcutaneous construct and abundant throughout the perivenous dermal matrix. Remodeled area was quantified as a percentage of the total dermal construct area, and nuclear density was quantified as nuclei/mm². Data are presented as mean ± SD (subcutaneous, n = 9; perivenous, n = 10). Statistical analysis was performed using separate two-tailed unpaired t-with Welch’s correction. Scale bars are indicated in the individual panels*.

In fibroblast-seeded constructs, beyond the vascular morphology, H&E analysis demonstrated greater dermal remodeling and cellularity following perivenous implantation. Remodeled area increased from 11.77 ± 2.82% in subcutaneous constructs to 17.73 ± 7.12% in perivenous constructs (p = 0.0311). Nuclear density similarly increased from 642.3 ± 127.5 nuclei/mm² to 832.9 ± 236.6 nuclei/mm² (p = 0.0437). Both comparisons were performed using two-tailed unpaired t-with Welch’s correction. These findings indicate that, in the presence of fibroblast seeding, perivenous implantation was associated with both greater structural remodeling and increased cellular repopulation of the dermal matrix.

Consistent with the micro-CT reconstructions, quantitative analysis confirmed significantly greater VV/TV in fibroblast-seeded perivenous constructs than in fibroblast-seeded subcutaneous constructs (Figure 9), increasing from 0.598 ± 0.189% to 7.329 ± 1.215% (p < 0.0001; two-tailed unpaired t-with Welch’s correction).

**Figure 9.**
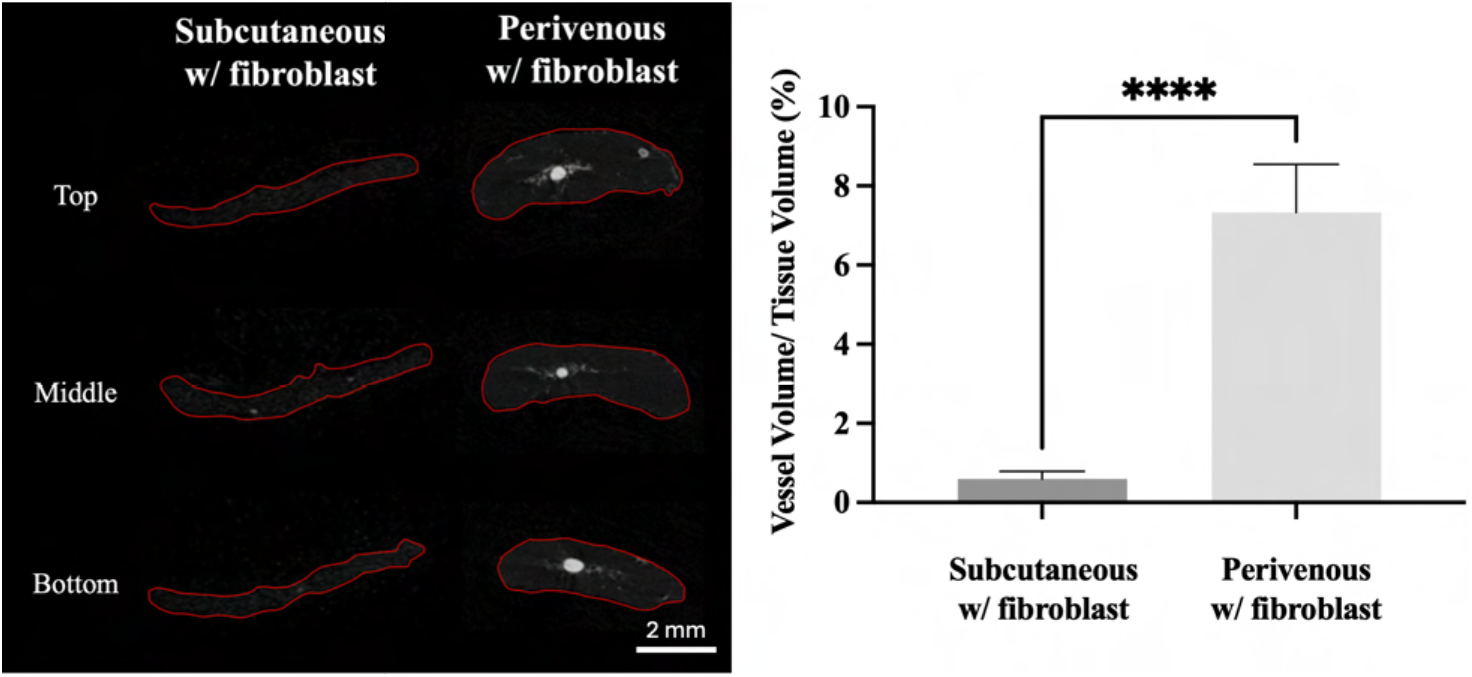
Perfused vascular volume in fibroblast-seeded decellularized dermal constructs after 8 weeks. Representative transverse micro-CT sections from the top, middle, and bottom of fibroblast-seeded subcutaneous and perivenous constructs are shown, with the dermal implant outlined in red. Vessel volume relative to tissue volume (VV/TV, %) was quantified within the dermal implant. The native femoral vein was excluded from the vascular-volume calculation in the perivenous group. Data are presented as mean ± SD (subcutaneous, n = 9; perivenous, n = 10). Statistical analysis was performed using a two-tailed unpaired t-test with Welch’s correction. ****p < 0.0001.

Finally, CD31 and α-SMA immunohistochemistry provided independent histological confirmation of this vascular difference (Figure 10). CD31-positive vessel density increased from 4.666 ± 2.738 vessels/mm² in subcutaneous constructs to 36.47 ± 11.61 vessels/mm² in perivenous constructs (p = 0.0008), while α-SMA-positive vessel density increased from 2.985 ± 1.908 to 33.51 ± 11.29 vessels/mm² (p = 0.0010 for both comparisons; two-tailed unpaired t-with Welch’s correction).

**Figure 10.**
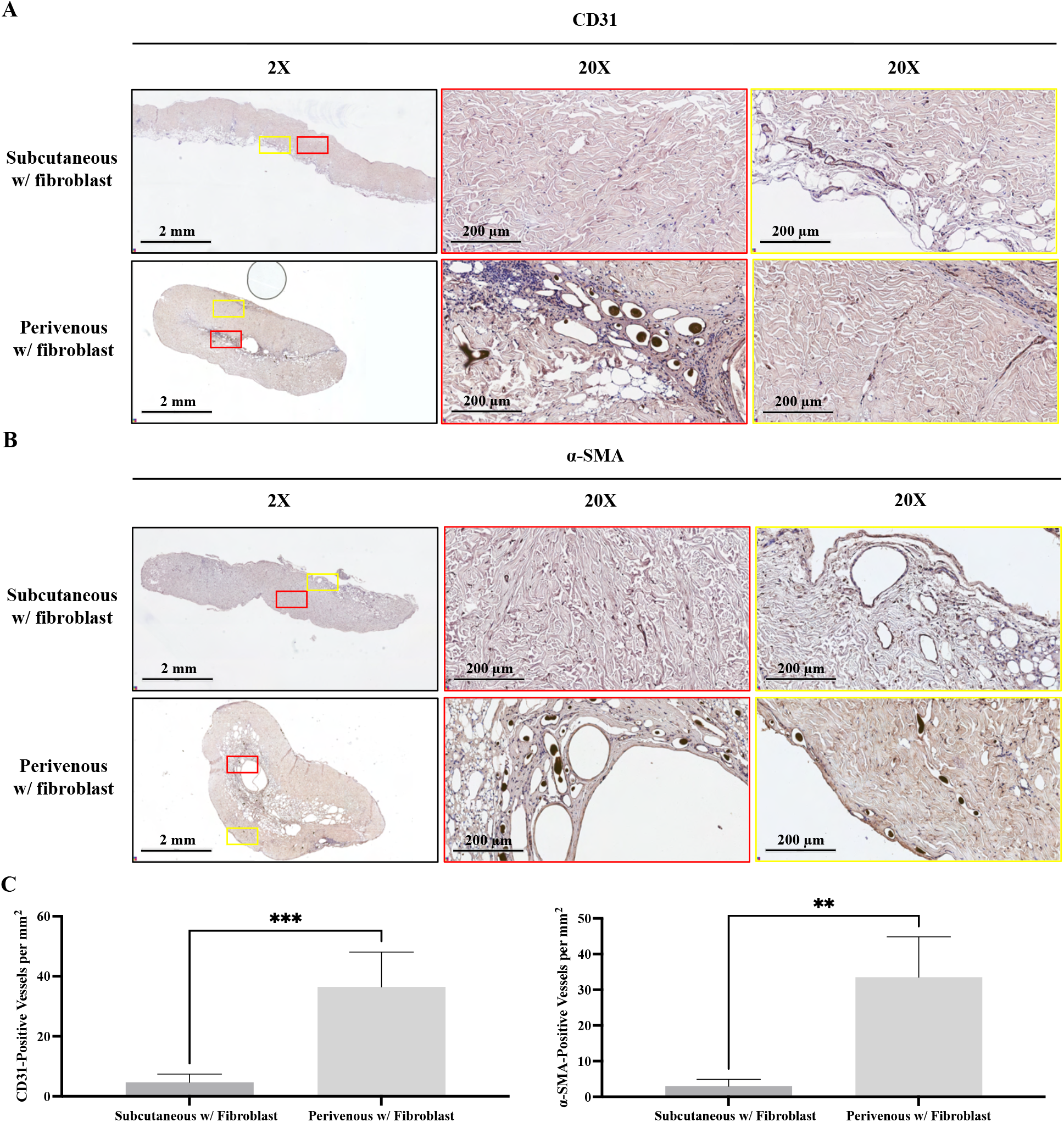
CD31-and α-SMA-positive vascularization of fibroblast-seeded decellularized dermal constructs after 8 weeks. Representative CD31 and α-SMA immunohistochemical staining of fibroblast-seeded subcutaneous and perivenous constructs is shown at low and high magnification, with corresponding regions indicated in the low-magnification images. CD31-positive and α-SMA-positive vessel densities were quantified as positive vascular structures per mm² of implant area. Data are presented as mean ± SD (n = 6 per group). *Statistical analysis was performed using separate two-tailed unpaired t-with Welch’s correction. **p < 0.01, ***p < 0.001. Scale bars are indicated in the individual panels*.

### 3.5. Integrated effects of implantation site and fibroblast seeding

To determine the independent and combined effects of implantation site and fibroblast seeding, the same 8-week vascular and histological data were further analyzed using two-way ANOVA (Figure 11). Implantation site significantly affected VV/TV, CD31-positive vessel density, α-SMA-positive vessel density, and nuclear density (all p < 0.0001), whereas fibroblast seeding significantly affected VV/TV (p < 0.0001), remodeled area (p < 0.0001), and nuclear density (p = 0.0039). A significant site × fibroblast interaction was detected only for VV/TV (p < 0.0001), indicating that the effect of fibroblast seeding on vascular volume depended on implantation site. Uncorrected Fisher’s LSD comparisons showed that fibroblast seeding increased VV/TV within the subcutaneous group (p < 0.0001), whereas the difference within the perivenous group was not significant (p = 0.0849). Fibroblast seeding also increased remodeled area within both implantation sites. No significant fibroblast effect was detected for CD31-or α-SMA-positive vessel density.

**Figure 11.**
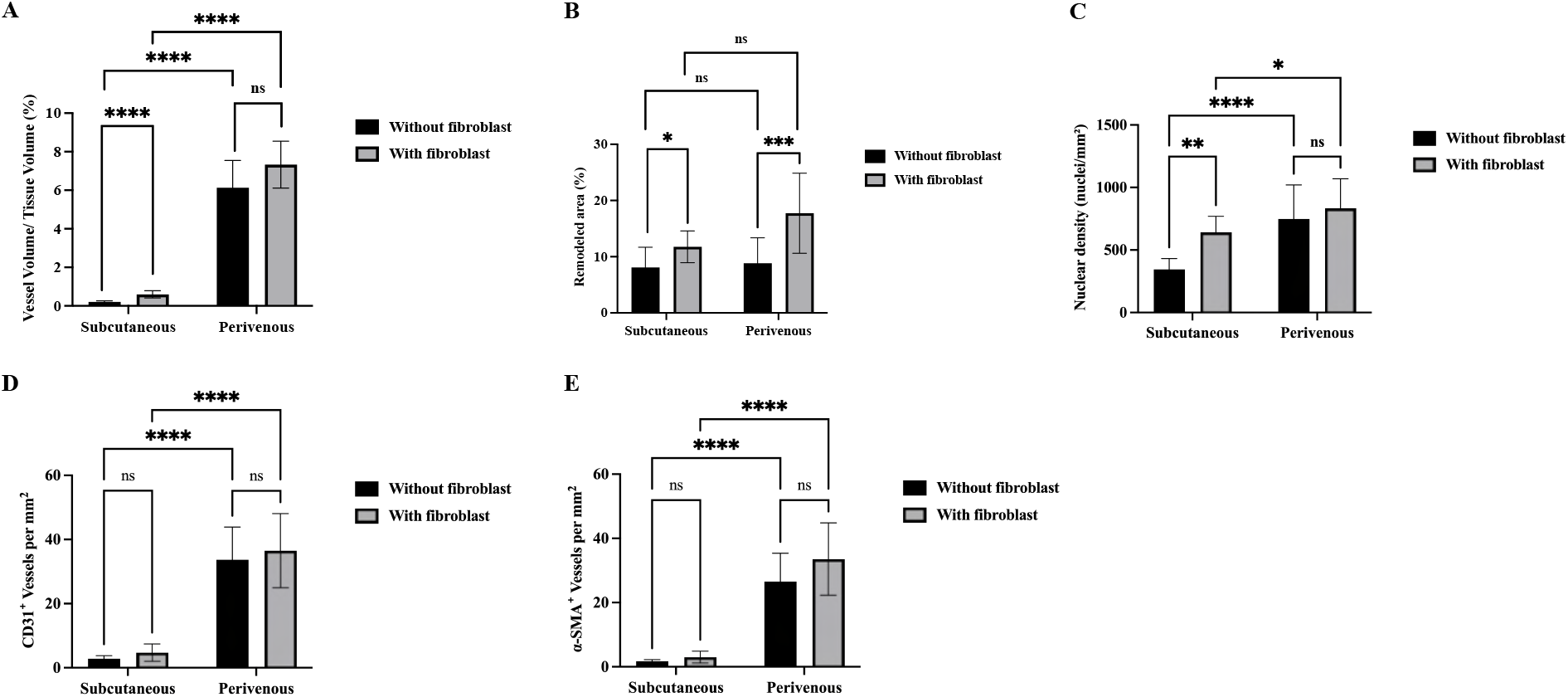
Integrated effects of implantation site and fibroblast seeding on vascularization and dermal remodeling after 8 weeks. (A) Vessel volume/tissue volume (VV/TV, %). (B) Remodeled area (%). (C) Nuclear density (nuclei/mm²). (D) CD31-positive vessel density (vessels/mm²). (E) α-SMA-positive vessel density (vessels/mm²). Data are presented as mean *± SD on the original scale. The same 8-week data presented in the preceding analyses were further analyzed using two-way ANOVA to assess the effects of implantation site (subcutaneous versus perivenous), fibroblast seeding (without versus with fibroblasts), and their interaction, followed by Uncorrected Fisher’s LSD comparisons for indicated pairwise comparisons. VV/TV, remodeled area, and α-SMA-positive vessel density were log-transformed for statistical analysis; nuclear density and CD31-positive vessel density were analyzed on the original scale. Reported p values correspond to the scale used for statistical analysis. ns, not significant; *p ≤ 0.05, **p ≤ 0.01, ***p ≤ 0.001, ****p ≤ 0.0001*.

### 3.6. Perivenous dermal constructs remained functionally perfused after mobilization

At 8 weeks, the perivenous dermal construct was surgically exposed and completely dissected free from the surrounding tissues while preserving only its connection to the femoral vein (Figure 12 and Supplementary Video S1). The femoral vein remained continuous within the mobilized construct with visible ongoing blood flow.

**Figure 12.**
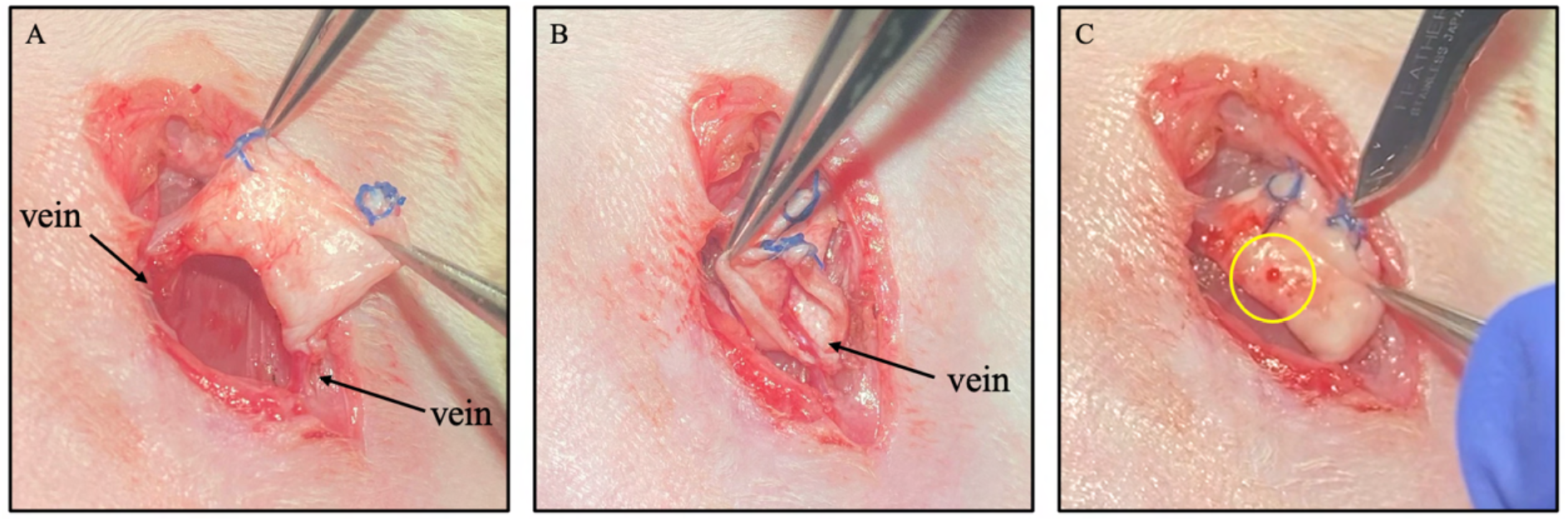
Functional perfusion of a perivenous decellularized dermal construct after 8 weeks. Representative intraoperative images showing exposure and mobilization of the dermal construct surrounding the femoral vein. (A) The femoral vein remained continuous within the construct following dissection from the surrounding tissues. (B) The femoral vein remained visibly patent with ongoing blood flow after construct mobilization. (C) Superficial incision of the dermal scaffold resulted in visible bleeding (circled in yellow), demonstrating functional perfusion of the vascularized construct. The corresponding procedure is shown in Supplementary Video S1.

Superficial incision of the mobilized dermal construct with a scalpel resulted in visible bleeding from the scaffold, demonstrating that blood continued to reach the intradermal vascular network through the remaining femoral venous connection. This observation complemented the Microfil-perfusion, micro-CT, and immunohistochemical findings and provided functional evidence that the vascular structures within the perivenous dermis were perfused.

### 3.7. Perivenous vascularization and fibroblast seeding improved healing after flap elevation

Following confirmation that the perivenous construct remained perfused after mobilization, the functional consequence of this vascularization was assessed after flap elevation. Wound healing after flap elevation differed markedly between the subcutaneous and perivenous constructs (Figure 13). In the subcutaneous groups, a dark eschar developed over the exposed dermal surface after flap elevation and persisted throughout the 28-day observation period, with wound area gradually decreasing. Fibroblast seeding significantly influenced wound-area reduction, with significant effects of time (p < 0.0001), fibroblast seeding (p = 0.0024), and the time × fibroblast interaction (p = 0.0189). Šídák-adjusted comparisons showed significantly lower wound area in fibroblast-seeded constructs at days 1 (p = 0.0078), 4 (p = 0.0114), and 28 (p = 0.0232). At day 28, the remaining wound area was 67.75% in the subcutaneous group without fibroblasts and 59.24% in the group with fibroblasts (Figure 13A,B).

**Figure 13.**
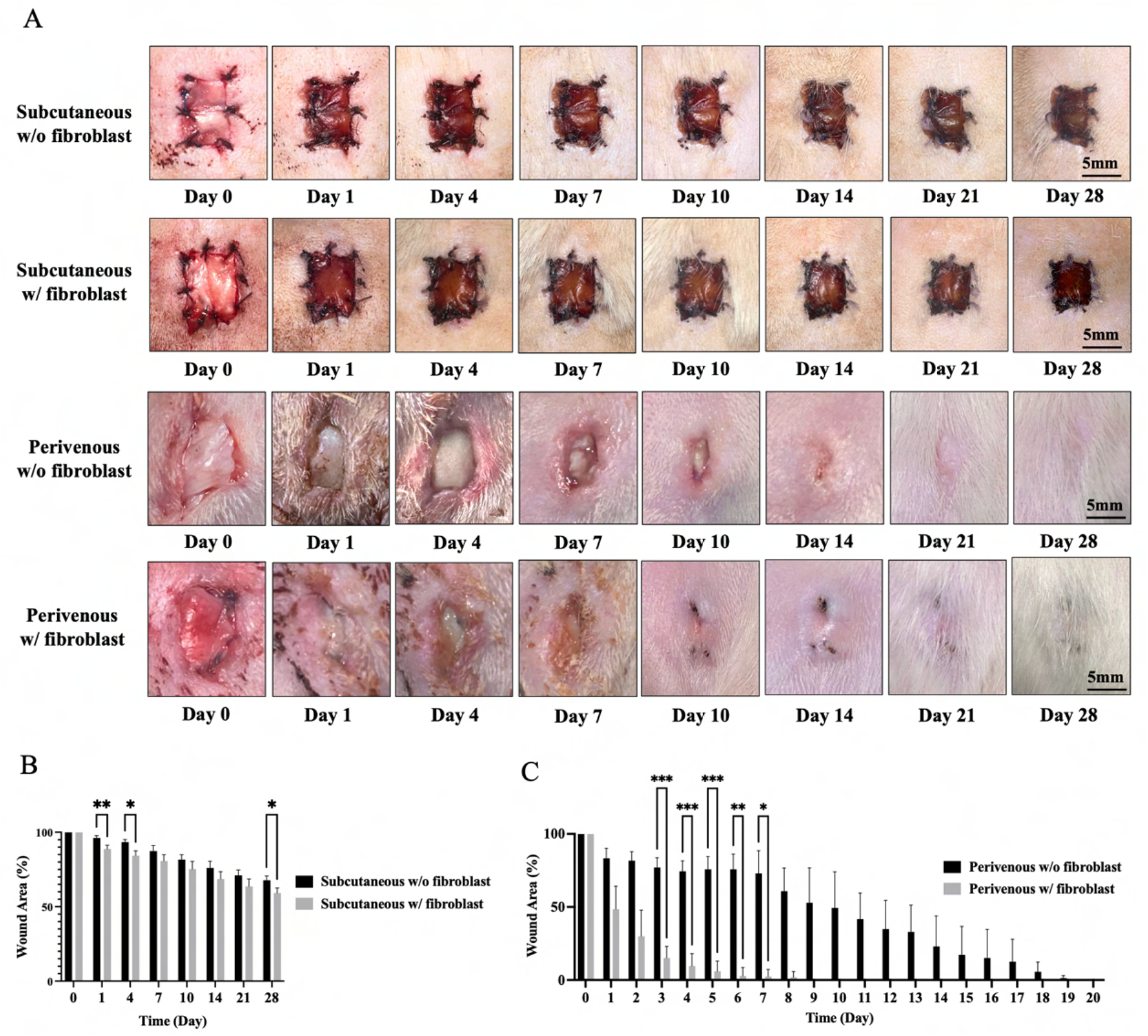
Wound healing following flap elevation of subcutaneous and perivenous dermal constructs. (A) Representative serial photographs of subcutaneous and perivenous constructs with and without fibroblasts from day 0 to day 28 after flap elevation. Scale bars: 5 mm. (B) Quantification of wound area over time in the subcutaneous groups. (C) Quantification of wound area over time in the perivenous groups. Data are presented as mean ± SD (subcutaneous, n = 5/group; perivenous, n = 4/group). Longitudinal wound-area data were analyzed using mixed-effects models with time as the repeated factor and fibroblast seeding as the between-subject factor, followed by Šídák’s multiple comparisons test. ns, not significant; *p ≤ 0.05; **p ≤ 0.01; ***p ≤ 0.001.

In contrast, the perivenous constructs showed progressive closure of the exposed dermal surface without the persistent dark eschar observed in the subcutaneous groups. Wound area decreased substantially faster following fibroblast seeding. Mixed-effects analysis showed significant effects of time, fibroblast seeding, and a time × fibroblast interaction (all p < 0.0001). Šídák-adjusted comparisons showed significantly lower wound area in the fibroblast-seeded group from days 3 to 7 (day 3, p = 0.0005; day 4, p = 0.0006; day 5, p = 0.0005; day 6, p = 0.0020; day 7, p = 0.0327), whereas differences at days 1-2 and from day 8 onward were not significant. The perivenous constructs with fibroblasts reached complete closure earlier than those without fibroblasts (Figure 13A,C).

### 3.8. DiI-associated fluorescence remained predominantly within the dermal implant after flap elevation

To determine where the seeded fibroblast-associated signal persisted during healing, for the two fibroblast-seeded groups, fluorescence microscopy was used to assess the distribution and persistence of DiI-associated fluorescence within the healed flap and residual dermal implant at day 28 after flap elevation (Figure 14).

**Figure 14.**
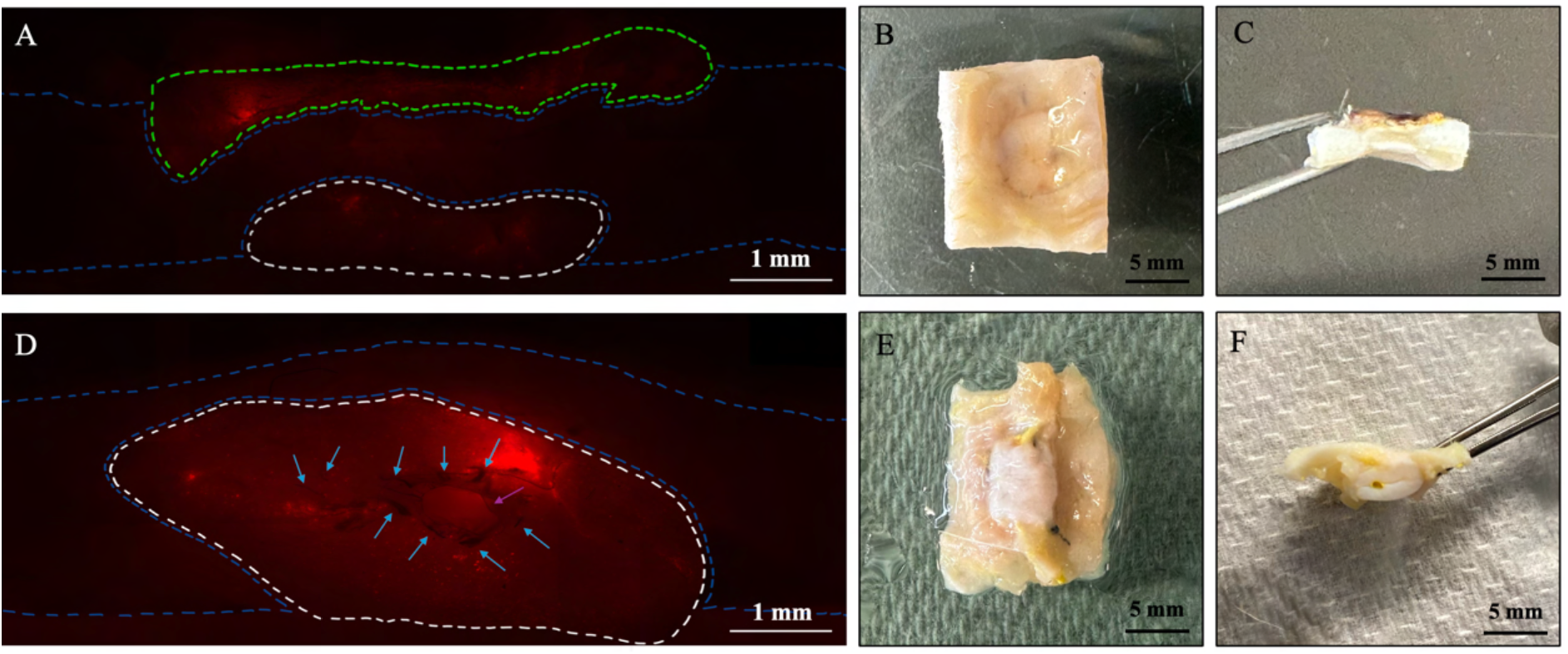
Fluorescence and gross imaging of fibroblast-seeded subcutaneous and perivenous constructs after flap elevation. *(A) Subcutaneous flap showing DiI-associated fluorescence within the eschar (green) and residual dermal implant (white). (B) Underside of the flap showing the residual implant. (C) Cross-section showing eschar, newly formed tissue, and dermal implant. (D) Perivenous flap showing the dermal implant around the femoral vein (purple arrow) and Microfil-perfused vessels (blue arrows). (E) Underside of the perivenous flap showing the vein-associated implant. (F) Cross-section showing the implant around the femoral vein beneath newly formed tissue. Scale bars = 1 mm (A,D) and 5 mm (B,C,E,F)*.

In the subcutaneous group, a dark eschar remained over the exposed surface of the construct. The residual dermal implant was clearly distinguishable beneath the newly formed tissue, although it appeared reduced compared with its original configuration (Figure 14B,C). Cross-sectional examination showed a layered architecture consisting of the surface eschar, intervening newly formed tissue, and residual dermal implant. Strong DiI fluorescence was detected predominantly within the residual dermal implant and the overlying eschar, whereas substantially weaker fluorescence was observed within the intervening newly formed tissue (Figure 14A). This distribution indicated that fibroblast-associated DiI signal remained primarily within or close to the original implanted material rather than being extensively distributed throughout the newly formed tissue.

In the perivenous group, no surface eschar was observed at day 28. Newly formed superficial tissue covered the dermal implant, and the implant remained associated with the femoral vein within the flap (Figure 14E,F). DiI fluorescence was concentrated predominantly within the residual dermal implant, particularly around the femoral vein, while little or no detectable DiI fluorescence was observed within the newly formed superficial tissue (Figure 14D). Microfil-filled vessels (blue arrows) were also visible within and around the vein-associated dermal implant, consistent with maintained perfusion of the vascularized construct. These findings indicate that fibroblast-associated DiI signal remained predominantly associated with the implanted dermal matrix after flap elevation, with limited detectable signal in the newly formed overlying tissue.

In the corresponding groups without fibroblast seeding, the gross and cross-sectional tissue architecture was comparable to that of their respective fibroblast-seeded groups, while no DiI fluorescence was detected.

### 3.9. Perivenous flaps showed tissue integration and organized skin architecture after elevation

Consistent with the gross healing and fluorescence findings, H&E staining at day 28 demonstrated distinct tissue organization between the subcutaneous and perivenous flaps (Figure 15). Consistent with the gross observations in Figure 13A, both subcutaneous groups retained a prominent surface eschar (green frames). Beneath the eschar, the residual dermal implant was surrounded by collagenous connective tissue, while a continuous organized epidermal layer and skin appendages were not clearly identified in the regions examined (yellow frames). At the margins of the construct, the implanted dermis showed continuity with the surrounding host connective tissue, demonstrating integration at the implant-host interface (blue frames). Cellular material was also present within portions of the eschar.

**Figure 15.**
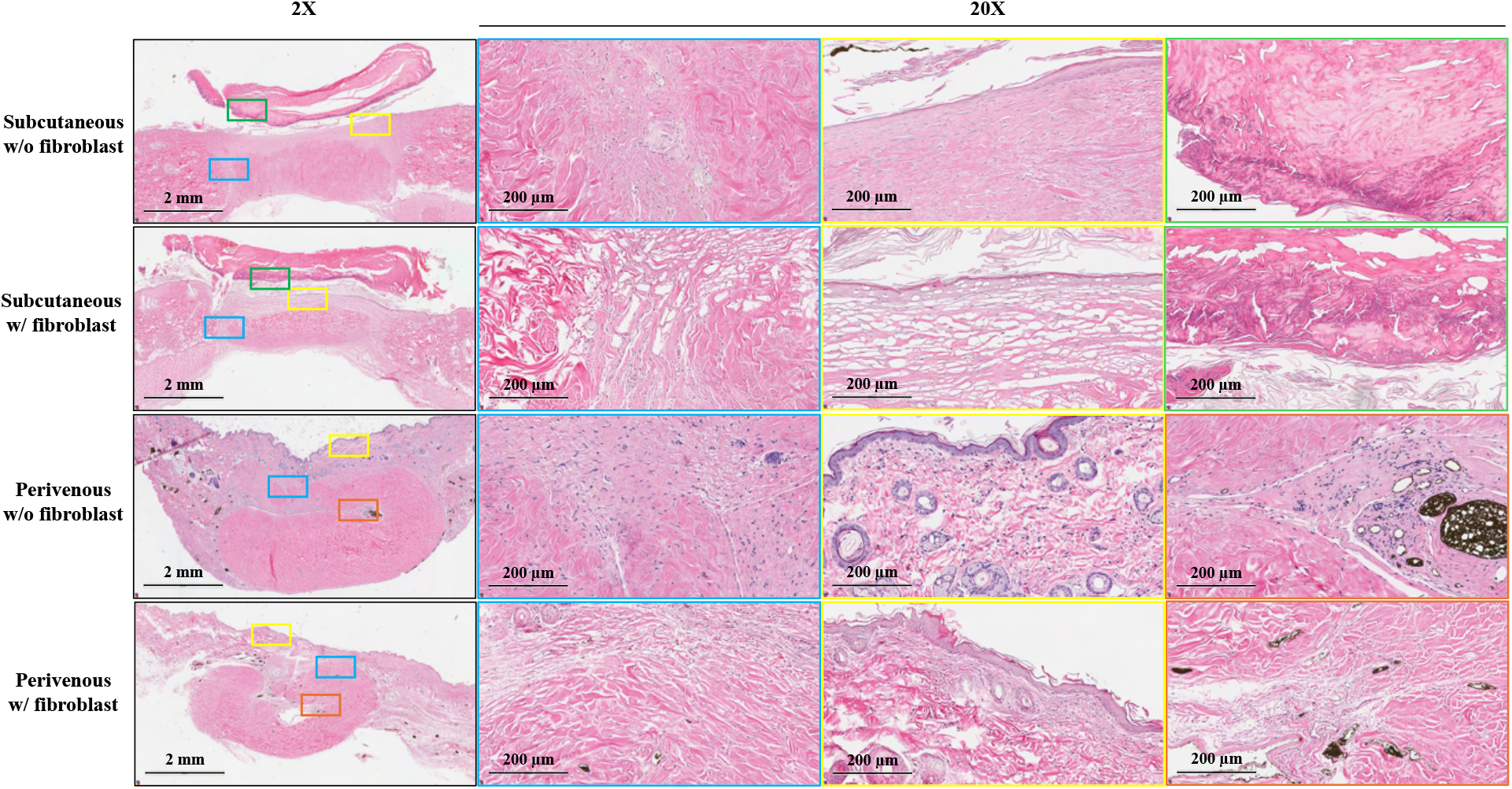
Histological organization of subcutaneous and perivenous dermal constructs after flap elevation. Representative H&E-stained sections at day 28 showing subcutaneous and perivenous constructs with and without fibroblast seeding. Blue frames show integration between the dermal implant and surrounding host tissue. Yellow frames highlight the superficial tissue, including hair follicles and sebaceous gland-like structures in the perivenous constructs. Green frames indicate the surface eschar in subcutaneous constructs. Orange frames show the vein-associated region with vascular profiles and Microfil-containing vessels. Scale bars = 2 mm (2×) and 200 µm (20×).

Consistent with this observation, DiI fluorescence remained detectable within the eschar of fibroblast-seeded subcutaneous constructs in Figure 14A, indicating persistence of DiI-associated fluorescence within the eschar at day 28.

The perivenous flaps showed a more organized superficial tissue architecture. The implanted dermis was closely integrated with the surrounding host tissue, with continuity of collagenous tissue across the implant-host interface (blue frames). The superficial tissue showed epithelial coverage together with hair follicles and associated sebaceous gland-like structures (yellow frames), consistent with the hair-bearing appearance observed macroscopically after flap elevation (Figure 13A). The vein-associated region remained identifiable within the construct, with vascular profiles and Microfil-containing vessels retained within the surrounding tissue (orange frames). Similar overall histological features were observed with and without fibroblast seeding. Together, these findings demonstrate integration of the implanted dermis with host tissue in both implantation conditions, while perivenous constructs showed more organized superficial tissue and preservation of the vein-associated vascularized region after flap elevation.

### 3.10. Perivenous flaps retained vascular networks after flap elevation

Finally, to determine whether the vascular differences observed histologically were reflected in the three-dimensional perfused network after elevation, micro-CT angiography was used to assess the three-dimensional vascular architecture of the flaps 4 weeks after elevation (Figure 16). In both subcutaneous groups, Microfil-perfused vessels were present throughout the surrounding host tissue but remained sparse within the outlined dermal implant region (Figure 16A,B). In contrast, the perivenous flaps retained a prominent vascular network within the construct region, with branching Microfil-perfused vessels distributed around the femoral vein and throughout the vein-associated dermal implant (Figure 16C,D).

**Figure 16.**
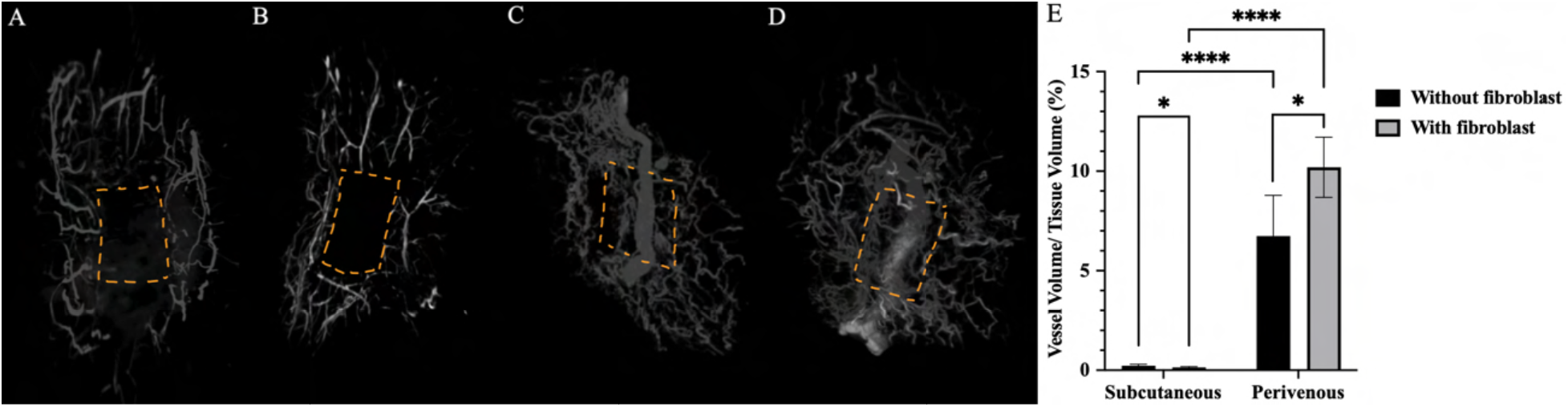
Three-dimensional vascular architecture of elevated dermal flaps. Micro-CT angiography of subcutaneous (A–B) and perivenous (C–D) constructs without (A, C) or with fibroblast seeding (B, D). Orange dashed outlines indicate the approximate dermal implant region. (E) Vessel volume relative to tissue volume (VV/TV, %) within the dermal implant region after flap elevation. Data are presented as mean ± SD on the original scale (subcutaneous: n = 5/group; perivenous: n = 4/group). VV/TV was log-transformed and analyzed using two-way ANOVA followed by uncorrected Fisher’s LSD comparisons test. *Reported p values correspond to the log-transformed analysis. ns, not significant; *p ≤ 0.05; ****p ≤ 0.0001*.

Quantitative analysis confirmed these differences (Figure 16E). Two-way ANOVA showed a significant effect of implantation site (p < 0.0001), no significant overall effect of fibroblast seeding (p = 0.8281), and a significant implantation site × fibroblast interaction (p = 0.0030). VV/TV was markedly greater in the perivenous groups than in the corresponding subcutaneous groups, both without and with fibroblast seeding (both p < 0.0001). Within the subcutaneous groups, fibroblast seeding decreased VV/TV from 0.225 ± 0.071% without fibroblasts to 0.136 ± 0.043% with fibroblasts (p = 0.0128). In contrast, fibroblast seeding increased VV/TV within the perivenous constructs from 6.742 ± 2.044% to 10.19 ± 1.510% (p = 0.0407).

Together, these findings demonstrate that the vascular advantage associated with perivenous implantation was retained 4 weeks after flap elevation. The significant interaction indicates that the effect of fibroblast seeding on post-elevation vascularization depended on implantation site, with opposite directional effects: VV/TV decreased with fibroblast seeding in subcutaneous constructs but increased with fibroblast seeding in perivenous constructs.

## 4. Discussion

The present study demonstrates that a single unmodified vein can support extensive vascularization of a compliant decellularized dermal matrix. Positioning the matrix around the femoral vein resulted in an approximately 30-fold increase in perfused vessel volume compared with conventional subcutaneous implantation, and this difference was supported by substantially greater CD31-and α-SMA-positive vessel densities. Importantly, the pre-existing femoral vein was excluded from the micro-CT vascular-volume calculation, such that the measured VV/TV represented the perfused vascular network associated with the dermal implant rather than the axial vein itself. These findings extend the material-induced venosome concept from rigid bioceramic scaffolds to a biologically derived soft-tissue extracellular matrix[27, 28]. Previous studies showed that the vascular response around a vein is strongly influenced by scaffold material, with monetite producing substantially greater vascularization than microporous polymeric scaffolds and different bioceramic compositions producing different responses[28, 29]. Decellularized dermis represents a distinct material environment, with a native extracellular-matrix architecture rather than an engineered porous structure, demonstrating that the single-vein response is not restricted to calcium-phosphate biomaterials[36, 37].

The present findings also extend the state of the art in pre-vascularization of engineered soft tissue by showing that a compliant dermal matrix can be vascularized around a single intact vein without creation of an AV shunt or incorporation of an arterial vessel. Previous AV loop and AV bundle studies have established that defined vascular carriers can generate axially vascularized soft-tissue constructs and, more recently, support free-flap transfer in large-animal models[18, 19, 25, 26]. In contrast to these approaches, the present strategy preserved the native flow-through femoral vein and avoided microsurgical creation of an arteriovenous circuit during the prefabrication stage. The ability of the construct to remain perfused after mobilization and flap elevation therefore represents an important intermediate step toward a transferable vascularized dermal substitute.

The comparison with subcutaneous implantation is particularly informative. Acellular dermal matrices can gradually become vascularized through vessel ingrowth from the surrounding host tissue during host incorporation[38–40]. Decellularized human dermis has also been shown to support cellular infiltration and angiogenesis during wound healing[41, 42].

Importantly, the organization of the dermal matrix itself can influence the distribution of newly formed capillary networks during healing[42, 43]. Despite this capacity, the subcutaneous constructs in the present study remained only sparsely vascularized after 8 weeks, whereas the matrix positioned around the femoral vein developed an extensive branching vascular network. The pronounced difference between the two implantation conditions supports a strong vascularization benefit associated with the perivenous strategy.

The mechanism by which an unmodified vein promotes vascularization of the surrounding dermal matrix remains unclear. The adventitial and perivascular compartments of large vessels are biologically active and contain fibroblasts, immune cells, progenitor populations, pericytes, and endothelial cells associated with the vasa vasorum, which can participate in vascular remodeling and neovascularization[44–47]. The adventitia has therefore been described as an active regulator of vessel-wall structure and remodeling rather than simply an outer supporting layer[45]. Vasa vasorum networks are themselves dynamic and capable of angiogenic expansion in response to changes in the local vascular environment[48–50]. In the present study, vessels were observed around the femoral vein and extending into the adjacent dermal matrix, consistent with a possible contribution from this perivascular compartment.

Together, these observations support the femoral vein and its surrounding perivascular environment as an important vascular axis for development of the intradermal network.

The immunohistochemical findings provide further information on the organization of the vascular network. The marked increase in CD31-positive vessel density was accompanied by a similarly pronounced increase in α-SMA-positive vessel density in the perivenous constructs. This indicates that the perivenous response involved not only a greater abundance of endothelial vascular structures but also substantially more α-SMA-positive vascular structures[51]. Pericytes and vascular smooth-muscle cells contribute to vessel development, stabilization, remodeling, and vascular function[52, 53]. However, because α-SMA can also be expressed by myofibroblasts, the α-SMA findings are interpreted as complementary evidence of vascular organization rather than as a direct measure of vessel maturity[54].

Fibroblast seeding provided an additional biological component to the construct, promoting matrix remodeling and cellular repopulation at 8 weeks, while its influence on perfused vascular volume depended on the implantation environment. Notably, these changes were not accompanied by significant overall effects on CD31-or α-SMA-positive vessel density, suggesting that the influence of fibroblast seeding on VV/TV was not simply reflected by changes in vessel density. The DiI findings complement this interpretation by showing persistence of fibroblast-associated signal after 8 weeks and a broader distribution through the perivenous matrix than in subcutaneous or in vitro constructs.

After flap elevation, a site-dependent fibroblast effect was again observed. Fibroblast seeding was associated with reduced VV/TV in subcutaneous constructs but increased VV/TV in perivenous constructs, consistent with the significant implantation site × fibroblast interaction. This opposing response suggests that the vascular effect of fibroblast seeding was strongly influenced by the surrounding implantation environment rather than representing a uniform pro-angiogenic effect. Dermal fibroblasts can support angiogenesis through paracrine signaling and extracellular-matrix remodeling, while VEGF-expressing dermal fibroblasts have also been shown to enhance angiogenesis and wound healing[32, 33, 55–58].

Beyond the quantitative and histological evidence of vascularization, the most important functional observation was that the vascularized dermal construct remained perfused after complete dissection from the surrounding tissues. At 8 weeks, the construct was mobilized while maintaining its connection to the femoral vein, and a small scalpel incision into the isolated dermis produced visible bleeding (Figure 12; Supplementary Video S1). This provided functional support for the Microfil, micro-CT, and histological findings by showing that the intradermal vascular network remained connected to the host circulation after removal of the surrounding tissue attachments. Quantitative post-elevation micro-CT further showed that the vascular advantage of perivenous implantation was retained 4 weeks after elevation, with VV/TV remaining markedly greater in both perivenous groups than in the corresponding subcutaneous groups (both p < 0.0001). After elevation, subcutaneous constructs showed very low VV/TV, whereas perivenous constructs retained substantial vascular volumes. These findings support maintenance of a functional vascularized construct around the preserved femoral venous axis after mobilization.

This functional connection is particularly relevant to soft-tissue reconstruction because the vascularized dermal construct remained organized around a native macroscopic vessel that could be directly identified and surgically manipulated. The present approach therefore differs from conventional vascularization from the surrounding recipient bed by generating a vascularized soft-tissue volume around a defined axial vessel. This principle is consistent with the original material-induced venosome study, in which a construct generated around a single vein remained patent following transplantation[28]. Extending this concept to decellularized dermis is important because the compliant matrix can be folded, sutured, and contoured for soft-tissue reconstruction.

The functional importance of this vascularization became particularly evident after flap elevation. Subcutaneous constructs developed a persistent dark eschar and remained incompletely healed at day 28, whereas perivenous constructs showed progressive surface closure and more organized superficial tissue. Histology at day 28 further showed epithelial coverage, hair follicles, and associated sebaceous gland-like structures in the perivenous flaps. Together with the retained post-elevation vascular network, these findings indicate that perivenous vascularization was the dominant determinant of construct performance after elevation. Fibroblast seeding also influenced tissue healing and remodeling. Fibroblast-seeded constructs generally showed improved wound closure, although the timing of this effect differed between the two implantation sites. At 8 weeks, fibroblast seeding also increased tissue remodeling and affected cellular repopulation of the constructs. In contrast, its effect on vascularization was more dependent on the implantation environment. Before flap elevation, fibroblast seeding increased VV/TV significantly in subcutaneous constructs but not in perivenous constructs. After flap elevation, the effects diverged further, with VV/TV decreasing in subcutaneous constructs but increasing in perivenous constructs following fibroblast seeding. These findings suggest that fibroblasts contributed to tissue remodeling and healing, but did not produce a uniform increase in vascularization across implantation environments. This is consistent with the established role of dermal fibroblasts in wound repair, where they contribute to extracellular-matrix remodeling and communicate with keratinocytes and endothelial cells through paracrine signaling[59–61]. Together, these findings suggest that fibroblast seeding supported tissue remodeling and wound healing, while its effect on vascularization depended on the implantation environment.

Several limitations should be considered. The initial vascularization phase was assessed at a single 8-week endpoint and therefore does not define the time course of vascular development before flap elevation. In addition, the subcutaneous and perivenous groups differed in anatomical recipient bed and surgical handling; therefore, the contribution of site-specific factors cannot be excluded, and a locally matched control would be required to isolate the specific contribution of the venous axis. More detailed characterization of the decellularized matrix, including quantitative residual-DNA and immunogenicity analyses, would also strengthen future studies, as the histological absence of visible nuclei alone does not establish complete decellularization. The anatomical origin of the newly formed vessels was not directly traced. Similarly, although DiI-associated fluorescence remained detectable, DiI labeling cannot definitively establish the long-term identity or viability of the originally seeded fibroblasts. The 8-week and post-elevation measurements were obtained from separate animal cohorts and therefore do not represent longitudinal measurements. In addition, although acute bleeding after mobilization supports a functional vascular connection at that time, the 4-week post-elevation assessment does not establish exclusive long-term dependence on the femoral vein because new vascular connections with surrounding tissues may have developed. Complete detachment and microsurgical re-anastomosis were not performed; therefore, the ability of the vascularized dermal construct to function as a transferable free flap remains to be established. Finally, only male rats were studied, limiting the generalizability of these findings across sex. These limitations define important directions for further investigation but do not alter the central finding that a functional vascular network can be generated within decellularized dermis around a single unmodified vein.

## 5. Conclusion

In conclusion, this study demonstrates that extensive and functionally connected vascularization of a decellularized dermal matrix can be generated around a single unmodified vein. The findings extend the material-induced venosome concept beyond rigid calcium-phosphate scaffolds and show that the response can occur within a compliant biological extracellular matrix. At 8 weeks, implantation site strongly influenced vascularization, while fibroblast seeding influenced dermal remodeling, nuclear density, and perfused vascular volume, with its effect on VV/TV depending on implantation site.

Importantly, after removal of the surrounding tissue attachments, the vascularized dermis showed acute perfusion while its femoral venous connection was preserved, and quantitative micro-CT confirmed that the vascular advantage of perivenous implantation was retained 4 weeks after flap elevation. These observations do not establish exclusive long-term dependence on the femoral vein because vascular connections with surrounding tissues may develop after inset. Fibroblast seeding also accelerated wound closure, while its effect on post-elevation vascular volume differed between implantation sites. Together, these findings support the formation of a mobilizable vascularized soft-tissue volume organized around a surgically accessible venous axis and provide a basis for future evaluation of single-vein vascularized dermis as a reconstructive tissue construct, including true microsurgical free-flap transfer.

## Notes

### Competing Interest Statement

The authors have declared no competing interest.

### Funding

- Natural Sciences and Engineering Research Council of Canada (NSERC)
- Canadian Institutes of Health Research (CIHR)

